# Contraction of the ROS scavenging enzyme glutathione *S*-transferase gene family in cetaceans

**DOI:** 10.1101/596395

**Authors:** Ran Tian, Inge Seim, Wenhua Ren, Shixia Xu, Guang Yang

**Affiliations:** Jiangsu Key Laboratory for Biodiversity and Biotechnology, College of Life Sciences, Nanjing Normal University, Nanjing, Jiangsu, 210046, China; Integrative Biology Laboratory, College of Life Sciences, Nanjing Normal University, Nanjing, Jiangsu, 210046, China; Comparative and Endocrine Biology Laboratory, Translational Research Institute-Institute of Health and Biomedical Innovation, School of Biomedical Sciences, Queensland University of Technology, Woolloongabba, Queensland, 4102, Australia

**Keywords:** glutathione transferase, GST, gene family, cetaceans, oxidative stress adaptation

## Abstract

Cetaceans are a group of marine mammals whose ancestors were adaptated for life on land. Life in an aquatic environment poses many challenges for air-breathing mammals. Diving marine mammals have adapted to rapid reoxygenation and reactive oxygen species (ROS)-mediated reperfusion injury. Here, we considered the evolution of the glutathione transferase (GST) gene family which has important roles in the detoxification of endogenously-derived ROS and environmental pollutants. We characterized the cytosolic GST gene family in 21 mammalian species; cetaceans, sirenians, pinnipeds, and their terrestrial relatives. All seven GST classes were identified, showing that GSTs are ubiquitous in mammals. Some GST genes are the product of lineage-specific duplications and losses, in line with a birth-and-death evolutionary model. We detected sites with signatures of positive selection that possibly influence GST structure and function, suggesting that adaptive evolution of GST genes is important for defending mammals from various types of noxious environmental compounds. We also found evidence for loss of alpha and mu GST subclass genes in cetacean lineages. Notably, cetaceans have retained a homolog of at least one of the genes *GSTA1*, *GSTA4*, and *GSTM1*; GSTs that are present in both the cytosol and mitochondria. The observed variation in number and selection pressure on GST genes suggest that the gene family structure is dynamic within cetaceans. Taken together, our results indicate that the cytosolic GST family in cetaceans reflects unique evolutionary dynamics related to oxygen-poor aquatic environments.

## INTRODUCTION

One of the classical examples of evolution is the return of terrestrial vertebrates to an aquatic environment – manifested by functional (secondary) adaptations in species whose ancestors departed an aquatic environment hundreds of millions of years earlier. The order Cetacea (whales, dolphins, and porpoises) is a model group of air-breathing marine mammals that transitioned to an aquatic lifestyle approximately 55 million years ago (Uhen 2007). The majority of cetaceans make shallow, short dives. This includes the common dolphin (*Delphinus delphis*) which usually dives down to 200 m and remains submerged for 5 minutes (Schreer and Kovacs 1997). Some cetaceans are capable of deep, long dives. For example, the sperm whale (*Physeter catodon*) can dive to a depth of 3,000 m and stay underwater for at least 138 minutes (Schreer and Kovacs 1997). Regardless of their diving abilities, all cetaceans face the tremendous challenge posed by a lack of oxygen during dives, so-called asphyxia (the integration of hypoxia, hypercapnia, and acidosis) (Elsner and Gooden 1983). In response, cetaceans have numerous adaptations of the respiratory system, such as improved oxygen delivery and storage in blood and muscle, as well as increased activity of glycolytic enzymes (Ramirez et al. 2007). Cardiovascular adaptations, including bradycardia (slowed heart rate) and peripheral vasoconstriction, also play a vital role in the conservation of oxygen in cetaceans. They augment or maintain blood flow to the central nervous system and heart but reduce flow in peripheral tissues such as kidney, liver, and skeletal muscle (ischemia). In terrestrial mammals reperfusion injury occurs when blood flow and oxygen delivery is restored to ischemic tissues, resulting in oxidative stress (Panneton 2013; Zenteno-Savín et al. 2011; Cantú-Medellín et al. 2011). Oxidative stress reflects an imbalance between the generation of reactive oxygen species (ROS) and the cell’s ability to detoxify reactive intermediates and repair damage (Birben et al. 2012). Oxidative stress can damage cells by lipid peroxidation and alteration of protein and nucleic acid structures (Apel and Hirt 2004).

The current evidence suggests that diving marine mammals and hibernating terrestrial mammals (e.g. ground-squirrels and some bats) have adapted to rapid reoxygenation and ROS-mediated reperfusion injury (Hermes-Lima et al. 2015). Oxidative damage is limited in cetaceans due to an intrinsic protection against ROS by scavenging enzymes and nonenzymatic antioxidants. Ceteceans have increased blood levels of the reduced form of glutathione (GSH), one of the most important nonenzymatic ROS scavengers (Wilhelm Filho et al. 2002; García-Castañeda et al. 2017). Similarily, the blood levels of vitamin E (α-tocopherol), which acts as a nonenzymatic antioxidant by protecting against peroxidation (Niwa 1999), is elevated in bottlenose dolphins (*Tursiops truncatus*) compared to its terrestrial sister taxa (Kasamatsu et al. 2009). ROS scavenging enzymes include glutathione peroxidase (GPX) which consumes hydrogen peroxide, glutathione reductase (GRS) which recycles glutathione from glutathione disulfide, superoxide dismutase (SOD) which scavenges superoxide radicals, and glutathione-*S*-transferase (GST) which catalyzes the conjugation of glutathione (Birben et al. 2012; Wilhelm Filho et al. 2002; Dröge 2002). Studies have revealed that antioxidant enzyme activies is higher in ceteceans compared to terrestrial mammals (Birben et al. 2012; Wilhelm Filho et al. 2002). Taken together, it is now recognized that cetaceans reduce reperfusion injury in various ways, however the genetic changes associated with these adaptations remain elusive.

Gene family innovation may enable species to adapt to novel or stressful environments (Kondrashov 2012). Here, we consider the ROS scavenging enzyme gene family glutathione transferase (GST; EC 2.5.1.18). A handful of studies suggest that GST gene gain/loss and positive selection facilitate the adaptation to changing environments (Ding et al. 2017; Low et al. 2007; Monticolo et al. 2017; Khan 2014; Liu et al. 2015). The GST family is abundant and widely distributed in vertebrates, plants, insects, and microbes (Board and Menon 2013). In addition to conjugating GSH with reactive electrophilic compounds, some GSTs can also deteoxify hydroperoxides (Sherratt and Hayes 2002). Mammalian GSTs have been divided into three structurally distinct superfamily classes with separate evolutionary origins (Board and Menon 2013): cytosolic, mitochondrial, and microsomal transferases. Cytosolic GSTs represent the largest class and consists of seven distinct subclasses: alpha (α; encoded by human chr 6), mu (μ; chr 1), theta (θ; chr 22), pi (π; chr 11), zeta (ζ; chr 14), sigma (σ; chr 4), and omega (ω; chr 10). The number of genes in each subclass varies across the phylogenetic tree. For example, human alpha (*GSTA1* to *GSTA5*) and mu (*GSTM1* to *GSTM5*) have five enzymes each, omega (*GSTO1* and *GSTO2*) two members each, and zeta (*GSTZ*) has only one member (Table S1). Members within each GST subclass shares greater than 40% amino acid sequence identity, while members of different subclasses share less than 25% identity (Wu and Dong 2012). Although diverse mammals have retained each cytosolic GSTs subclass, there is variation in the number of genes in each subclass. For example, there is one pi subclass gene (*GSTP*) in humans and two pi subclass genes in mice (Bammler et al. 1994). In this study, we performed a comparative genomics analysis of cytosolic GST genes in 21 mammals, including seven cetaceans, to improve our knowledge on the genetic and evolutionary dynamics of the GST superfamily in mammals.

## MATERIALS AND METHODS

### Sequence retrieval

We obtained known human GST genes by perusing research articles and recent reviews (Board and Menon 2013; Morel et al. 2002) and by downloading coding sequences (CDS) from GenBank (http://www.ncbi.nlm.nih.gov) (Benson et al. 2018). GenBank accession numbers are listed in Table S1. Employing human protein sequences as queries, BLASTn searches were performed using an in-house Python script (see Supplemental Material, file 2) on local databases constructed from downloaded genomic sequences from 20 species. These included seven cetaceans: bottlenose dolphin (*Tursiops truncatus*), killer whale (*Orcinus orca*), Yangtze river dolphin (*Lipotes vexillifer*), Yangtze finless porpoise (*Neophocaena asiaeorientalis asiaeorientalis*), minke whale (*Balaena acutorostrata*), bowhead whale (*Balaena mysticetus*), and sperm whale (*Physeter macrocephalus*); two pinnipeds: Weddell seal (*Leptonychotes weddellii*), Pacific walrus (*Odobenus rosmarus divergens*); one sirenian: Florida manatee (*Trichechus manatus latirostris*); and ten terrestrial mammals: cow (*Bos taurus*), Tibetan yak (*Bos mutus*), sheep (*Ovis aries*), Tibetan antelope (*Pantholops hodgsonii*), dog (*Canis lupus familiaris*), horse (*Equus caballus*), a bat (little brown bat; *Myotis lucifugus*), mouse (*Mus musculus*), naked mole rat (*Heterocephalus glaber*), and (African) elephant (*Loxodonta africana*). The completeness of the annotated gene set of the 20 specices was assessed using Benchmarking Universal Single-Copy Orthologs (BUSCO v3.0) with mammalian-specific single-copy orthologs (mammalia_odb9) (Simão et al. 2015). Genome sequencing and assembly information of each species are listed in Table S2.

To identify GST genes, we set the BLASTn parameter *E*-value cut-off to 10 and hits with the highest score, lowest *E*-value, and a length of ≥ 150bp were retained. We next retrieved multiple non-redundant hits by extending 1,000 bp at both 5’ and 3’ ends to find exon boundaries following the canonical gt/ag exon/intron junction rule (Cheng et al. 1995). We compared the genomic locations of each coding sequence among all genes to filter out repeated sequences with the same location on the same scaffold. The online resource GENEWISE (http://www.ebi.ac.uk/Tools/psa/genewise) (Birney et al. 2004) was employed to identify open reading frame (ORF) for all obtained sequences. Finally, all the predicted GST sequences were verified by BLAST against its respective species genomes to acquire complete GST gene sets. Mammalian GSTs were named according to the human GST nomenclature (Nebert and Vasiliou 2004). All identified GST genes were categorized into three categories – based on amino acid composition, unique motifs, BLAST and alignment results: 1) intact gene, a complete CDS region with the canonical structure typical of GST families; 2) partial gene, putative functional protein, but missing a start codon and/or stop codon; 3) pseudogene, highly similar to functional orthologs but with (a) inactivating mutation(s) and/or stop codon(s). To achieve a high accuracy in identifying GST genes in mammals, we used Genomicus v93.01 (Nguyen et al. 2017) to identify genes flanking the GST gene clusters in human and searched the mammalian genomes using BLAST to identify orthologous genomic regions. This enabled the identification of the correct arrangement and orientation of the GST genes in each species.

### Phylogenetic analyses

The phylogenetic relationships between putative GST members in each subclass were estimated by maximum likelihood and Bayesian methods, as implemented in RAxML v8.0.26 (Stamatakis 2014) and MrBayes v3.1.2 (Ronquist and Huelsenbeck 2003). Nucleotide sequences were aligned using MUSCLE v3.6 (Edgar 2004), implemented in SeaView v4.5.4 (Gouy et al. 2009), and manually corrected upon inspection. MrModeltest was used to estimate the best-fit model of nucleotide substitution (SYM+G) and amino acid substitution (JTT+G4) (Nylander 2009). For the RAxML analyses the ML phylogeny was estimated with 1,000 bootstrap replicates. For MrBayes analyses we performed two simultaneous independent runs for 50 million iterations of a Markov Chain, with six simultaneous chains and sampling every 1,000 generations. A consensus tree was obtained after discarding the first 25% trees as burn-in.

### Gene family analyses

To identify expanding and contracting gene ortholog groups across the mammalian phylogeny, we estimated the gene numbers on internal branches using a random birth and death process model implemented in the software CAFÉ v3.0, a tool for the statistical analysis of the evolution of the size of gene families (De Bie et al. 2006). We defined gene gain and loss by comparing cluster size differences between ancestors and each terminal branch among the phylogenetic tree. An ultrametric tree, based on the concatenated orthogroups, was estimated with BEAST v1.10 using Markov chain Monte Carlo (MCMC) with fossil calibrations and a Yule tree prior (Suchard et al. 2018). The molecular timescales were obtained from TimeTree (http://www.timetree.org) (Kumar et al. 2017). The analyses ran for 10 million generations, with a sample frequency of 1,000 and a burn-in of 10%.

### Adaptive evolution analyses

To evaluate the positive selection of all GST genes during mammalian evolution, codon substitution models implemented in the *codeml* program in PAML v4.4 (Yang 2007) were applied to GST gene alignments. Two pairs of site-specific modes were compared using the likelihood ratio test (LRT): M8 (beta & ω) vs. M8a (beta & ω = 1) (Swanson et al. 2003). M8 estimates the beta-distribution for ω and takes into account positively selected sites (ω > 1), with the neutral model M8a not ‘allowing’ a site with ω > 1. We next employed branch-site models (test 2) to explicitly assess the rate of evolution on a site along a specific lineage of a tree: branch-site model (Ma) vs. branch-site model with fixed ω1 = 1 (Ma0) (Zhang et al. 2005). The Ma model assumes that sites in the foreground branch (i.e. the branch of interest) are under positive selection. When the LRT was significant (*p* < 0.05) under the M8 and Ma tests, codon sites under positive selection were assessed using the Bayes Empirical Bayes (BEB) method (Yang et al. 2005). The species tree was used as the guide tree in all analyses (Figure S1). Multiple testing for positive selection on genes was corrected by performing a false discovery rate (FDR, Benjamini-Hochberg) test at a cutoff of 0.05 (Benjamini and Hochberg 1995). We only accepted positively selected sites with a posterior probability (PP) > 0.80. A series of models implemented in HyPhy (http://www.datamonkey.org) (Pond and Muse 2005) were also used to estimate ratios of nonsynonymous (*d_N_*) and synonymous (*d_S_*) based on a maximum likelihood (ML) framework (Pond and Frost 2005a). These models tested were Single Likelihood Ancestor Counting (SLAC), Fixed Effect Likelihood (FEL), and Random Effect Likelihood (REL) (Pond and Frost 2005b). As a criteria to identify candidates under selection, we used a Bayes Factor > 50 for REL and *P-*value of 0.10 for SLAC and FEL. The program TreeSAAP (Selection on Amino Acid Properties using Phylogenetic trees) v3.2 (Steve et al. 2003) was used to evaluate amino acid residue replacement during evolution, taking into account physicochemical properties and the assumption of random replacement under a neutral model of evolution. TreeSAAP complements *d_N_*/*d_S_*analysis, which does not distinguish between different types of non-synonymous substitutions. InterProScan 5 was used to annotate positively selected sites found in functional protein domains (Philip et al. 2014). We also used the PyMOL (http://pymol.sourceforge.net/) to load and manipulate PDB files, highlighting the position of selected residues.

### Comparison of mammalian niches

To investigate the potential links between the molecular evolution of GST genes and ecological adaptations, we assigned habitats (aquatic vs. terrestrial) according to data in the literature (Uhen 2007). We used Clade Model C (CmC) to identify the level of divergent selection among clade with different ecological niches and to test what partition of clades best fit the data (Bielawski and Yang 2004). CmC, which allows site variation among a priori defined foreground (cetacean, pinnipeds, sirenians) and background partitions (Figure S1), was compared with the M2a_rel null mode which does not allow variation among divisions. Taking into account the phylogenetic relationships between mammals, a phylogenetic ANOVA (Garland et al. 1993) using the function *phylANOVA* implemented in the R package ‘phytools’ (Revell 2012) was performed to test for differences in the number of GST genes between marine and terrestrial mammals.

## RESULTS

### Cytosolic glutathione transferase gene repertoires in mammals

To examine the evolution of cytosolic GST genes in mammals, we interrogated public genomic data of the 21 species, representing all major mammalian taxa. We identified a total of 448 GST genes (333 intact genes, 22 partial genes, and 93 pseudogenes) (Table 1). Amino acid sequence identity between GST paralogous was more than 50%, whereas the identity between all subclasses was less than 30% (Table S3). It has been reported that a low-quality genome can effect gene family analyses by introducing frame shifting errors in coding sequences (Young et al. 2010). In our study we employed genomes with good reported genome assemblies in an effort to minimize this potential bias. In agreement, BUSCO analysis on the protein gene sets showed that most, except for bowhead whale (74.6%) and Weddell seal (87.3%), included >95% complete sequences of mammalian universal single-copy orthologs (*n* = 4,104) (Table S2). This suggests that the genome assembly quality is similar between the marine and the terrestrial species.

**Table 1.** Overview of cytosolic glutathione transferase (GST) genes in 21 mammals.

| Subclass | Alpha<br>(GSTA) | Mu<br>(GSTM) | Theta<br>(GSTT) | Pi<br>(GSTP) | Zeta<br>(GSTZ) | Omega<br>(GSTO) | Sigma<br>(HPGDS) | Total number<br>of GSTs | Pseudogene<br>proportion |
| --- | --- | --- | --- | --- | --- | --- | --- | --- | --- |
| **Human** | 12(7:5:0) | 5(0:5:0) | 3(0:3:0) | 1(0:1:0) | 1(0:1:0) | 2(0:2:0) | 1(0:1:0) | 25(7:18:0) | 0.28 |
| **Mouse** | 6(1:5:0) | 10(4:6:0) | 3(0:3:0) | 7(0:7:0) | 1(0:1:0) | 2(0:2:0) | 1(0:1:0) | 30(5:25:0) | 0.17 |
| **Naked mole rat** | 4(1:3:0) | 10(0:10:0) | 2(0:2:0) | 2(0:2:0) | 1(0:1:0) | 2(1:1:0) | 1(0:1:0) | 22(2:20:0) | 0.09 |
| **Bottlenose dolphin** | 3(1:0:2) | 5(3:1:1) | 2(1:1:0) | 3(1:0:2) | 1(0:0:1) | 2(0:1:1) | 1(0:1:0) | 17(6:4:7) | 0.35 |
| **Killer whale** | 3(1:2:0) | 5(4:1:0) | 2(0:2:0) | 3(1:2:0) | 1(0:1:0) | 2(0:2:0) | 1(0:1:0) | 17(6:11:0) | 0.35 |
| **Yangtze finless porpoise** | 3(1:2:0) | 5(4:1:0) | 2(1:1:0) | 3(1:0:2) | 1(0:0:1) | 2(0:2:0) | 1(0:1:0) | 17(7:7:3) | 0.41 |
| **Yangtze river dolphin** | 3(1:2:0) | 5(4:1:0) | 2(1:1:0) | 3(1:2:0) | 1(0:1:0) | 2(0:2:0) | 1(0:1:0) | 17(7:10:0) | 0.41 |
| **Sperm whale** | 3(1:2:0) | 5(4:1:0) | 2(0:1:1) | 3(1:2:0) | 1(0:1:0) | 2(0:2:0) | 1(0:1:0) | 17(6:10:1) | 0.35 |
| **Minke whale** | 3(1:2:0) | 5(3:1:1) | 2(1:1:0) | 3(1:2:0) | 1(0:1:0) | 2(0:1:1) | 1(0:1:0) | 17(6:9:2) | 0.35 |
| **Bowhead whale** | 2(0:2:0) | 5(3:1:1) | 2(1:1:0) | 3(1:0:2) | 1(0:0:1) | 2(0:2:0) | 1(0:1:0) | 16(5:7:4) | 0.31 |
| **Cow** | 6(2:4:0) | 8(2:6:0) | 2(0:2:0) | 2(0:2:0) | 1(0:1:0) | 3(0:3:0) | 1(0:1:0) | 23(4:19:0) | 0.17 |
| **Tibetan yak** | 6(1:5:0) | 6(0:6:0) | 3(0:3:0) | 2(0:2:0) | 1(0:1:0) | 3(0:3:0) | 1(0:1:0) | 22(1:21:0) | 0.05 |
| **Sheep** | 7(1:6:0) | 8(2:6:0) | 2(0:2:0) | 2(0:2:0) | 1(0:1:0) | 4(0:4:0) | 1(0:1:0) | 25(3:22:0) | 0.12 |
| **Tibetan antelope** | 6(1:5:0) | 6(0:6:0) | 2(0:2:0) | 2(0:2:0) | 1(0:1:0) | 3(0:3:0) | 1(0:1:0) | 21(1:20:0) | 0.05 |
| **Weddell seal** | 4(1:2:1) | 5(2:3:0) | 2(0:2:0) | 7(1:4:2) | 1(0:1:0) | 2(0:2:0) | 1(0:1:0) | 22(4:15:3) | 0.18 |
| **Pacific walrus** | 5(1:4:0) | 8(2:6:0) | 2(0:2:0) | 6(1:5:0) | 1(0:1:0) | 2(0:2:0) | 1(0:1:0) | 25(4:21:0) | 0.16 |
| **Dog** | 6(1:5:0) | 5(3:2:0) | 2(0:2:0) | 12(6:6:0) | 1(0:1:0) | 2(0:2:0) | 1(0:1:0) | 29(10:19:0) | 0.34 |
| **Horse** | 10(1:9:0) | 6(2:4:0) | 2(0:2:0) | 2(0:2:0) | 1(0:1:0) | 2(0:2:0) | 1(0:1:0) | 24(3:21:0) | 0.13 |
| **Microbat** | 6(0:6:0) | 5(1:3:1) | 4(0:4:0) | 6(5:1:0) | 1(0:1:0) | 2(0:2:0) | 1(0:1:0) | 25(6:18:1) | 0.24 |
| **Florida manatee** | 4(0:4:0) | 8(0:8:0) | 2(0:2:0) | 1(0:1:0) | 1(0:1:0) | 2(0:2:0) | 1(0:1:0) | 19(0:19:0) | 0.00 |
| **Elephant** | 3(0:3:0) | 8(0:8:0) | 2(0:1:1) | 1(0:1:0) | 1(0:1:0) | 2(0:2:0) | 1(0:1:0) | 18(0:17:1) | 0.00 |

The number outside brackets is the number of genes in a subclass, while the numbers in the brackets, separated by a colon, indicate the number of pseudogenes, intact gene, and partial genes. The orders of cetacean are shadowed.

### The phylogenetic relationships of cytosolic GST genes

We next wished to classify the 448 cytosolic GST genes into their respective subclass. Phylogenetic relationships were reconstructed using maximum likelihood and Bayesian methods. Both topologies yielded similar branch patterns, indicating a reliable tree structure (Figure S2). Each GST subclass was clustered into a monophyletic group with high node bootstrap values (94-100% of bootstrapping) (Figure S2). However, subtrees of species within each subclass were not clearly resolved (< 50% node bootstrap support) (Figure S2) – especially for the alpha and mu subclasses where duplication events were common. According to the phylogenic tree, the cytosolic GST gene family is highly conserved in mammals (Figure S2). We detected 105 alpha subclass (81 genes and 24 pseudogenes), 133 mu subclass (90 genes and 43 pseudogenes), 47 theta subclass (42 genes and 5 pseudogenes), 74 pi (54 genes and 20 pseudogenes), 21 zeta subclass (21 genes and 0 pseudogenes), 47 omega subclass (46 genes and 1 pseudogenes), and 21 sigma subclass (21 genes and 0 pseudogenes) genes in the 21 species examined.

Notably, ML and Bayesian phylogenies based on the 448 GST genes did not arrange the seven subclasses into well-supported clades (< 50% node bootstrap support). This could result from the 93 pseudogenes and 22 partial GST genes that are likely no longer under natural selection pressure. Therefore, only the 333 intact (complete CDS) sequences were used to infer the phylogenetic tree and determine the evolutionary relationship of subclasses. The tree generated using an amino acid substitution model (Figure S3) was consistent with the tree structure obtained using nucleotide sequences (Figure 1). According to the new phylogeny, the mu subclass clustered with the pi subclass but the bootstrap support value and posterior probability were low (BS = 18%; PP = 0.34, Figure 1). The alpha subclass grouped with sigma with high bootstrapping and posterior probability (BS = 91%; PP = 0.58, Figure 1). The mu, pi, alpha, and sigma subclasses were much more closely related to each other, with 98% bootstrap support and 0.50 posterior probability (Figure 1). The theta subclass was placed as sister to a clade containing mu, pi, alpha, and sigma genes with high support (BS = 85%; PP = 1.00, Figure 1). Moreover, these five subclasses formed a monophyletic sister clade to the zeta subclass. The phylogenetic tree recovered the omega subclass as the most diverged lineage within GSTs (Figure 1).

**Figure 1.**
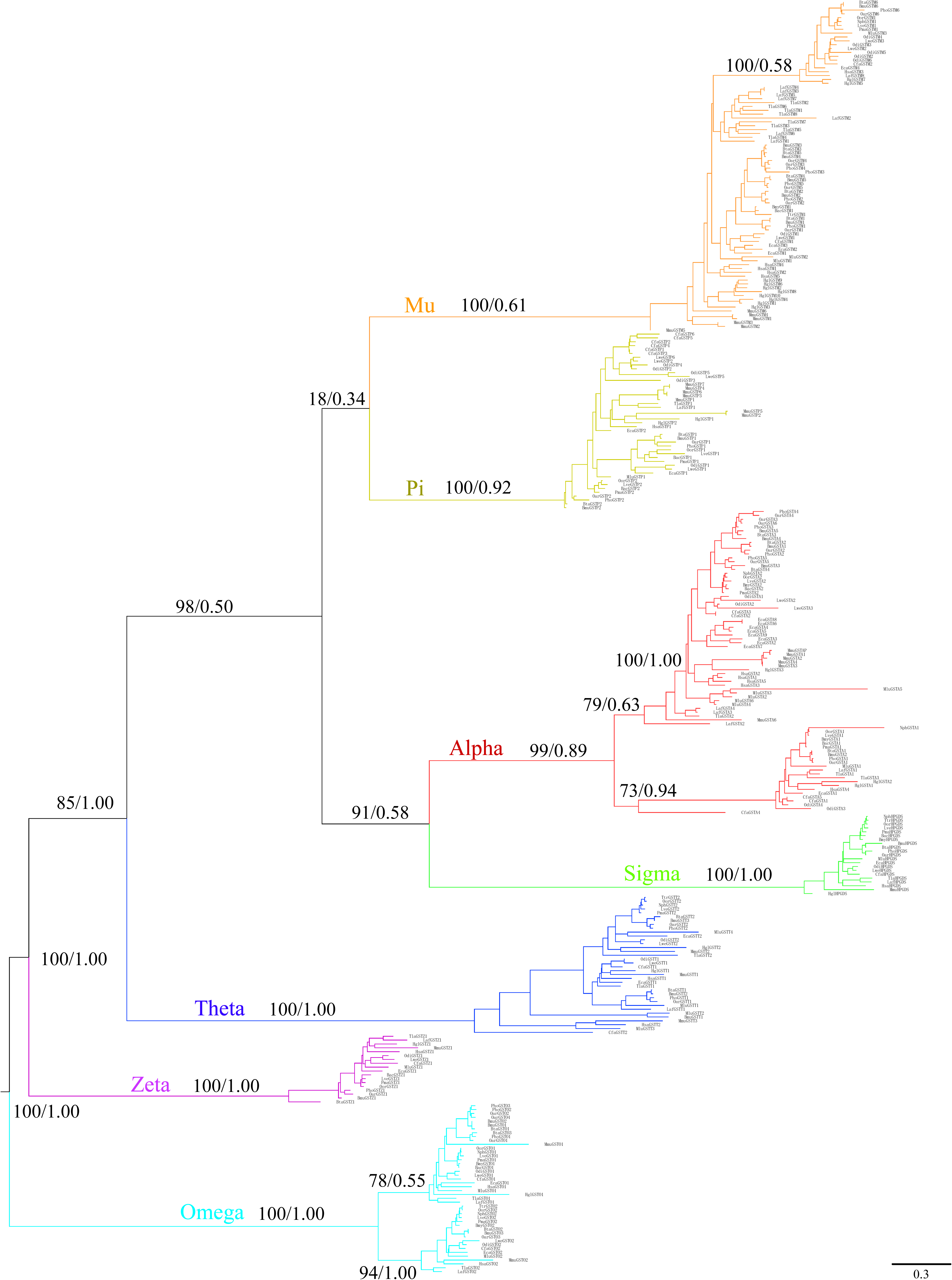
Phylogenetic tree of GST gene family in mammals. Maximum likelihood and Bayesian phylograms describing phylogenetic relationships among 333 intact (complete coding sequences) mammalian GSTs (21 species). Numbers on nodes correspond to maximum likelihood bootstrap support values and Bayesian posterior probabilities. The GST subclasses are indicated by different colours: alpha (red), mu (orange), pi (turquoise), omega (azure), sigma (green), zeta (purple), and theta (blue).

Further strengthening our GST gene predictions, all seven cytosolic GST subclasses showed conserved synteny and similar arrangement across mammalian genomes (Figure 2). Duplicated genes within each subclass were arranged in a tandem cluster, with two or more gene copies in tandem in the subclasses alpha, mu, theta, pi, and omega. The zeta and sigma classes had a single copy per species. The subclasses were flanked by the same pair of genes in all species (Figure 2), with the exception of the dog where pi subclass genes were present on separate scaffolds and possibly reflecting a sequencing or genome assembly artifact.

**Figure 2.**
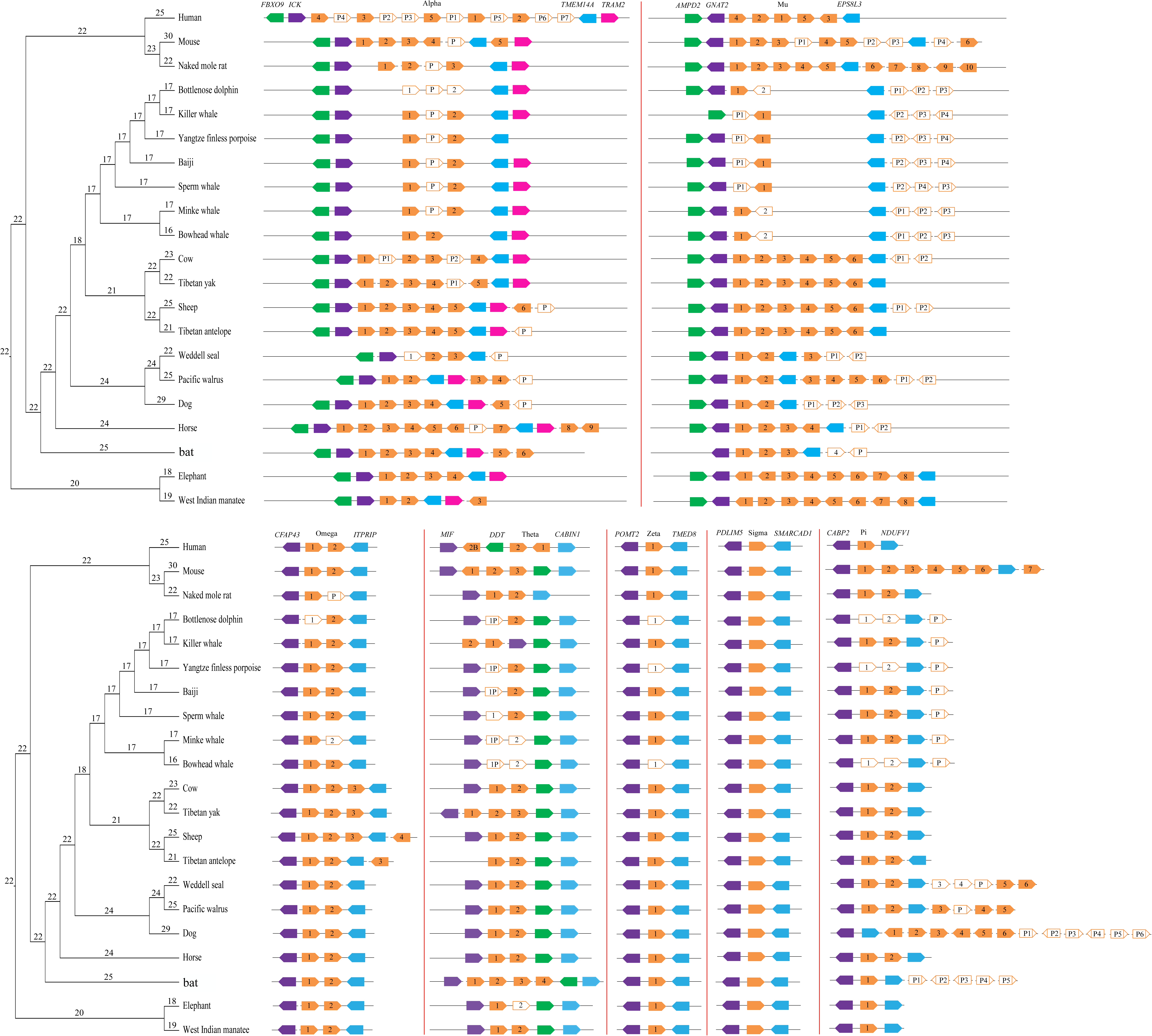
Genomic organization of GST genes in 21 mammalian species. The arrowed boxes represent genes and directions of transcription. GST genes are shown in orange, while flanking genes are indicated in green, purple, blue, and pink. Filled boxes: intact genes; empty boxes: partial gene, and empty boxes with a vertical line: pseudogenes (P). Connecting horizontal lines indicate genes on the same chromosome/genomic scaffold. Gene family sizes for ancestral states are shown along each node in the phylogenetic tree.

### Lineage-specific gene duplications and deletions

We next contrasted the number of GST gene copy number in mammalian lineages. Of the 21 species studied, mouse (25 intact GSTs), naked mole rat (20 intact GSTs), Tibetan yak (21 intact GSTs), Tibetan antelope (20 intact GSTs), sheep (22 intact GSTs), Pacific walrus (21 intact GSTs), and horse (21 intact GSTs) have the largest GST repertoires (Table 1, Figure 2). On the opposite spectrum, cetaceans appear to have the smallest GST repertoire – about ten functional GSTs per species (Table 1, Figure 2). In agreement, the fraction of GST pseudogenes is the highest in cetaceans (mean, 36%, Table 1), which is three times higher than terrestrial artiodactyls (mean, 11%, Table 1). Considering the alpha subclass, we found that the cetacean alpha subclass consists of two functional GSTs, but their relatives (i.e., artiodactyls) have at least four intact alpha GSTs. Sheep (6 intact GSTAs), horse (9 intact GSTAs), and bat (6 intact GSTAs) have a relatively large number of alpha GSTs. Similarly, only two functional mu GSTs were identified in cetaceans. In contrast, the number of functional GSTM genes in artiodactyls (6) is almost six times that of cetaceans. Notably, a large group of species, including all rodents and two afrotherians (West Indian manatee and African elephant), harbor six to ten functional mu GSTs (Table 1, Figure 2). Additionally, bat (4 intact GSTTs) has the largest gene number of theta GSTs, while the lowest gene copy number was found in cetaceans (just one gene in six cetaceans; two in killer whale). Almost all mammalian species have two omega subclass genes, with the exception of artiodactyls (3 or 4 genes) (Table 1, Figure 2). The zeta and sigma subclass gene number appear to be more conserved in mammals, with one copy identified, respectively. Furthermore, the gene gain and loss of GSTs at each ancestral node was estimated by the software CAFÉ (De Bie et al. 2006). We found that the terrestrial groups have similar GST gene repertoires – carnivorans (mean 24.8), rodents (mean 25), artiodactyls (mean 22.29) (Welch’s *t*-test, *p* > 0.05). In contrast, the number of intact GST genes in cetaceans (mean 16.92) was significantly lower than that of the terrestrial groups (Welch’s *t*-test, *p* < 0.001) (Figure 2). To compare gene numbers and account for statistical non-independence of closely-related species, we performed phylogenetic ANOVA (phylANOVA). After accounting for phylogeny, habitat (aquatic vs. terrestrial) was a signficant predictor of the gene copy number of all cytosolic GSTs combined (phylANOVA; *F* = 23.135, *p* = 0.009) (Figure 3A) and alpha-class GSTs (GSTA) alone (phylANOVA; *F* = 21.599, *p* = 0.007), with a smaller number of genes apparent in cetaceans (Figure 3B). In agreement, when we compared each marine order (Cetacea, Pinnipedia, Sirenia) to terrestrial species separately, cytosolic gene copy numbers was only significantly different between cetacean and terrestrial species (Figure 3C). This included all cytosolic GST combined (phylANOVA; *F* = 142.458, *p* = 0.001), GSTA alone (phylANOVA; *F* = 24.300, *p* = 0.026), and GSTM alone (phylANOVA; *F* = 27.557, *p* = 0.011).

**Figure 3.**
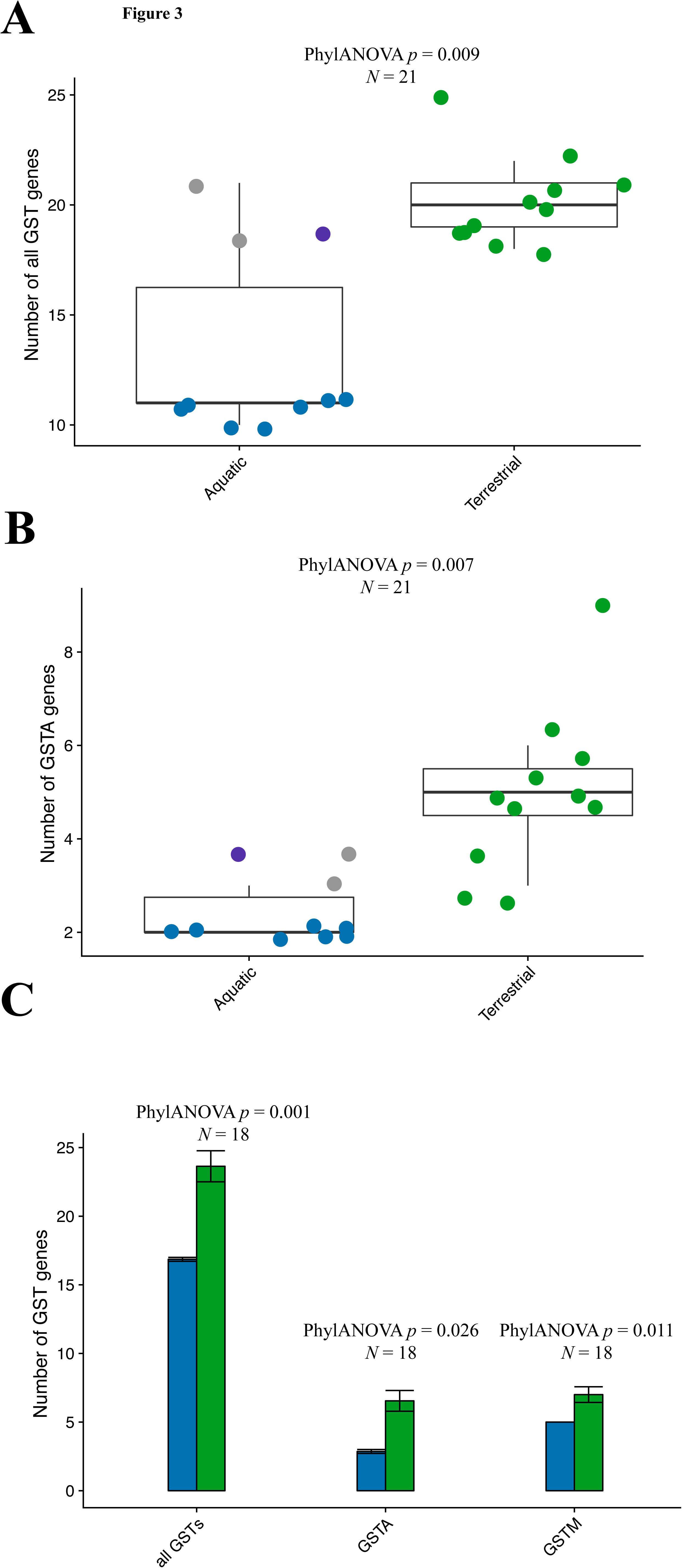
Differences in cytosolic GST genes between mammals inhabing aquatic and terrestrial habitats. (A) All GSTs genes combined, comparing aquatic and terrestrial mammals. Green dots indicate terrestrial species; blue dots, cetacean species; grey dots, pinnipeds; purple dots, sirenians. (B) alpha-class GSTs (GSTA) genes, comparing aquatic and terrestrial mammals. Annotated as in (A). (C) Comparison of all GST genes, alpha-class GSTs (GSTA) genes, and mu-class GSTs (GSTM) genes in ceteaceans and terrestrial mammals. Bar chart shows mean±s.e.m. Blue bars indicates cetacean species; green bars, terrestrial species. We compared each pair of distributions by phylogenetic ANOVA (phylANOVA) tests, which control for shared ancestry. The *p*-value for each test is shown above each plot.

### Evolutionary model of the cytosolic GST gene family in mammals

To investigate the possible role of natural selection on the evolutionary history of the cytosolic GST gene family, a series of site-specific and branch-specific evolutionary models were evaluated. Site-specific selection tests implemented in PAML (Yang 2007) were performed to assess the selective pressure acting on mammalian GSTs. Site-specific positive selection with posterior probabilities > 0.80 were detected for *GSTA1* (16 sites), *GSTM1* (5 sites), *GSTO1* (14 sites), *GSTO2* (3 sites), *GSTP1* (2 sites), *GSTP2* (6 sites), *GSTT2* (1 sites), and *GSTZ1* (3 sites) (Table S4); suggesting that GSTs have evolved under diversifying selection in mammals. Similarly, positively selected codons in eight genes were identified using SLAC (23 codons), FEL (28), and REL (27) models implemented in Datamonkey (Pond and Muse 2005). In total, 38 codons from eight genes (*GSTA1*: 10; *GSTM1*: 1; *GSTO1*: 9; *GSTO2*: 5; *GSTP1*: 2; *GSTP2*: 5; *GSTT2*: 3; and *GSTZ1*: 3) with evidence of positive selection were detected with at least two of the ML methods (Table 2). Among these codons, 31 sites were also detected by amino-acid level selection analyses using TreeSAAP (Steve et al. 2003). Intriguingly, site enrichment analysis revealed that the sites under positive selection (28/38, 73%) are located or close to a GSH binding site, a substrate binding pocket, a C-terminal domain interface, or a N-terminal domain interface (Table S5, Figure S4).

**Table 2.**
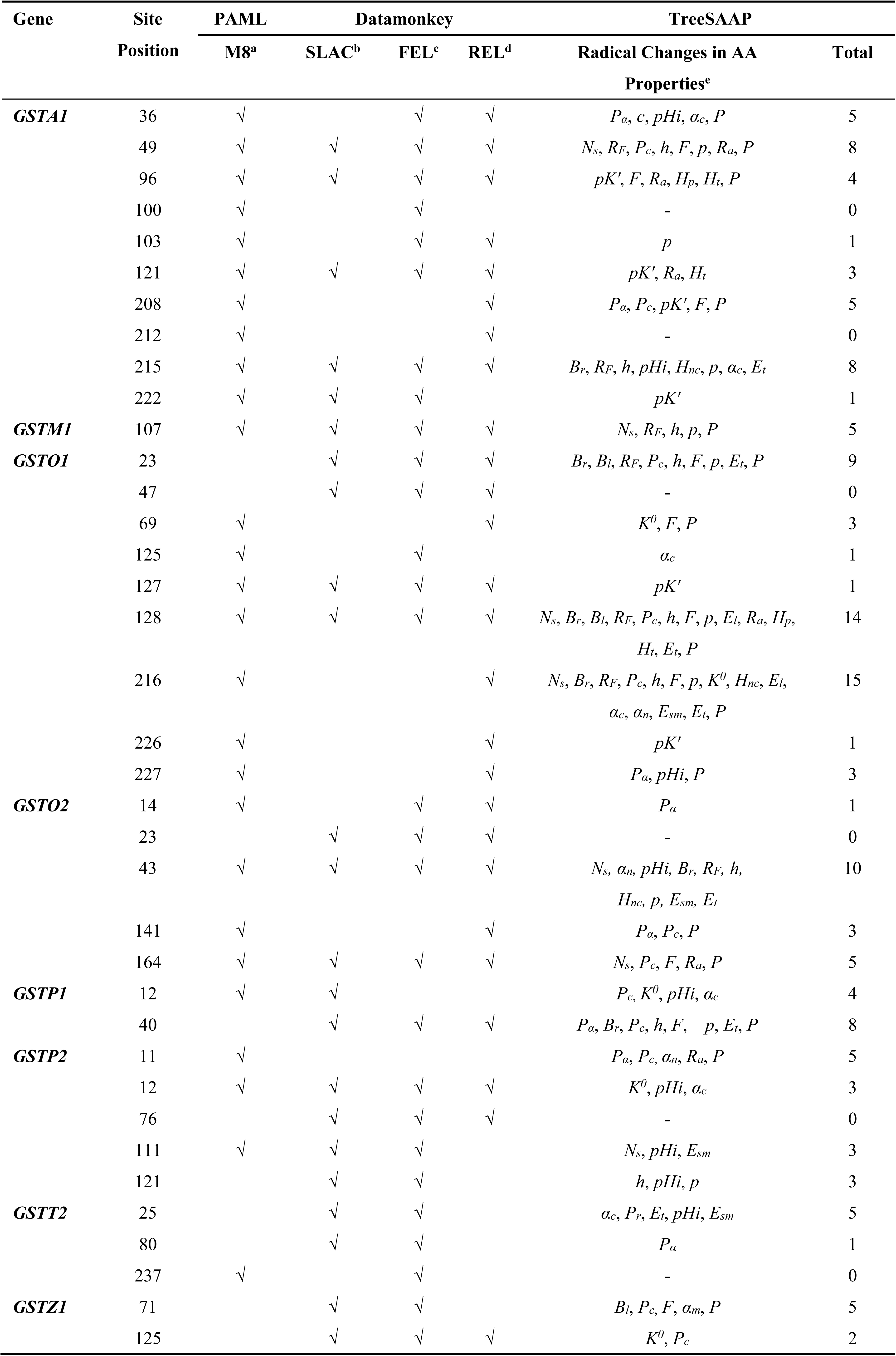
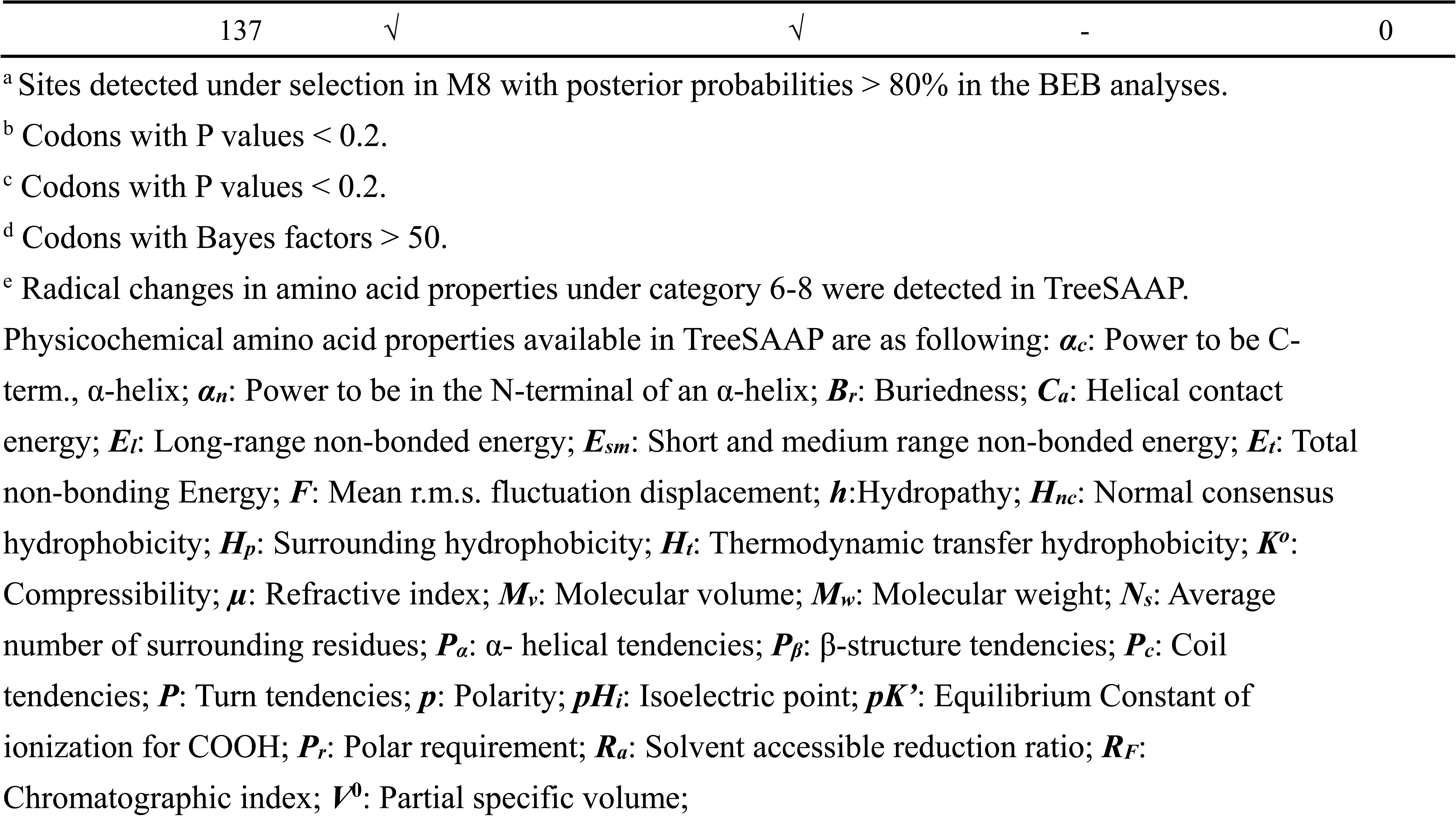
Amino acid sites under positive selection detected by ML methods

We also employed the PAML branch-site model (Yang 2007) to identify episodic adaptations that affect amino acids along specific lineages. Few internal branches and several terminal branches showed evidence of positive selection after FDR correction (Table 3, Figure S5). In cetaceans the lineage leading to *GSTO1* in bottlenose dolphin and *GSTP2* in sperm whale were under selection. In pinnipeds the branches leading to *GSTM1* in Pacific walrus and its ancestor, *GSTP2* in the ancestor, as well as *GSTA1* in Weddell seal were under positive selection. *GSTA1* and *GSTM1* were under positive selection only in pinnipeds (Pacific walrus and Weddell seal). The lineage leading to human *GSTA4*, elephant and bat *GSTT1*, cow, horse and manatee *GSTT2*, sheep and antelope *GSTP2*, naked mole rat *GSTO1*, as well as cetartiodactyla (includes whales and dolphins, and even-toed ungulates) *HPGDS* were also under positive selection.

**Table 3.**
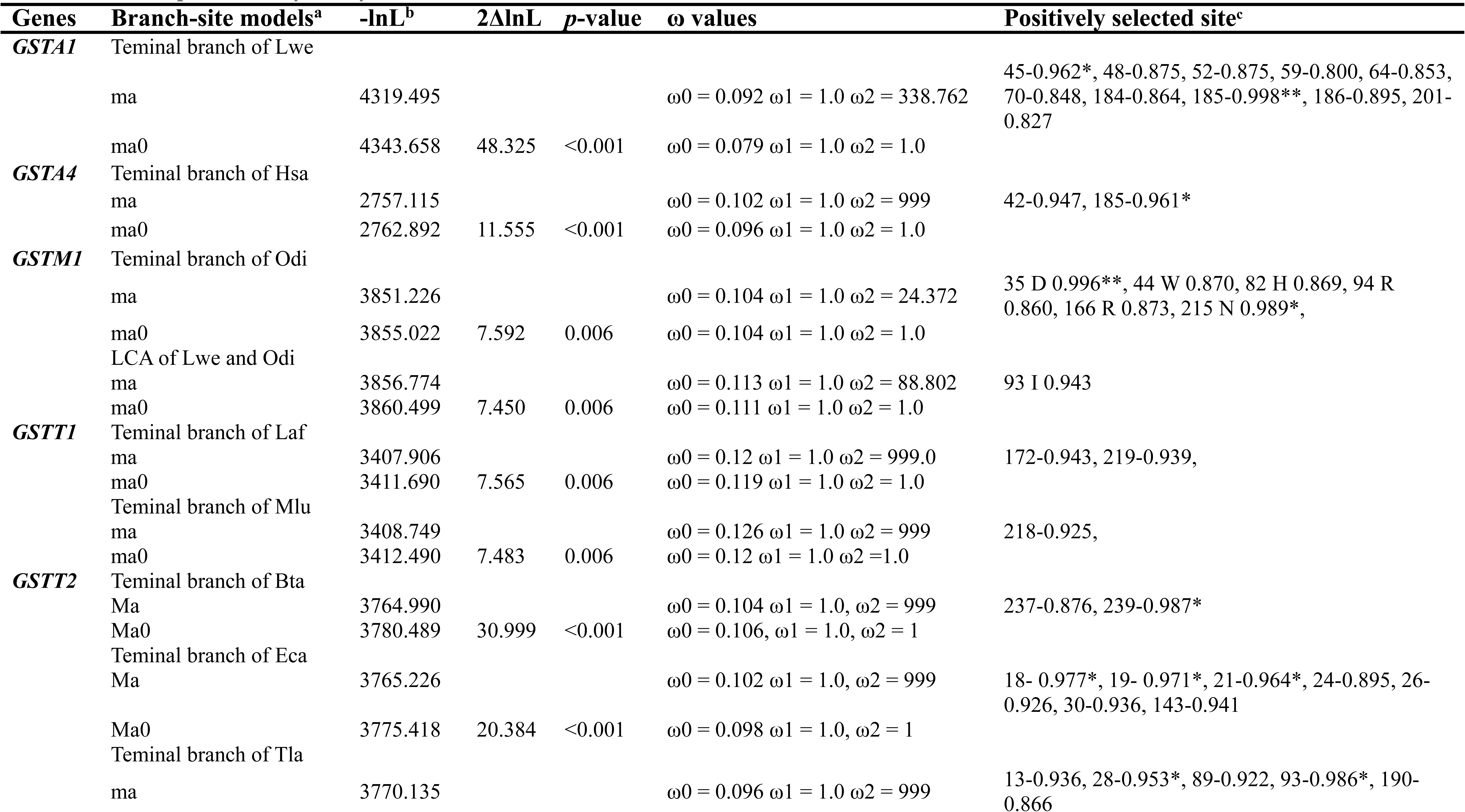
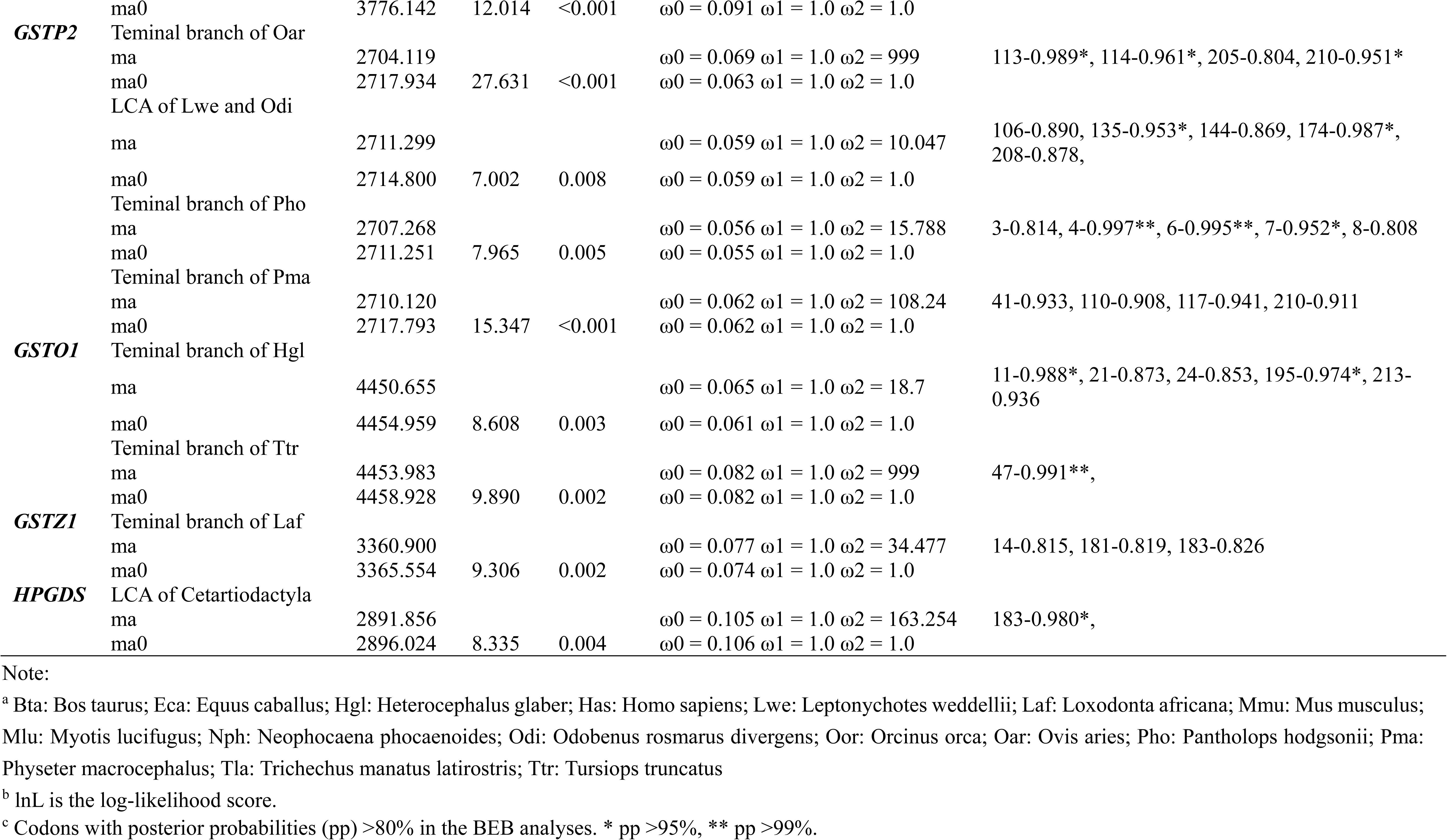
Selective pattern analyzed by Branch-site model

A more thorough investigation of the evolutionary history of mammalian GSTs was conducted by extending the analysis to comparing marine and terrestrial species. Clade model C allows for more than two clades to be defined as separate partitions, estimating ω separately for each partition. A model which assumes divergent selection along the cetacean, pinniped, sirenian, and marine mammal branches fitted the data better than the null model in the case of the GST genes *GSTA1*, *GSTA4*, *GSTM1*, *GSTM3*, and *GSTT1* (*p* < 0.05, Table S6), suggesting there is a divergent selection pressure between marine and terrestrial mammals and within marine mammal groups.

## DISCUSSION

### Molecular evolution of GST genes

Several previous surveys have documented the distribution of GST genes. Nebert and Vasiliou (2004) provided a comprehensive assessment of GST superfamily genes. Other studies examined the phylogeny (Pearson 2005) and evolution (da Fonseca et al. 2010) of the superfamily across the tree of life. In the present study, we expanded on the previous surveys by performing a comprehensive search for cytosolic GST genes in 21 mammals representative of all major mammalian taxa. Of these, 15 species had no previous information on their cytosolic GST gene repertoire (Figure 2). We provide evidence for positive selection acting on several GST genes in divergent taxonomic groups (*GSTA1*, *GSTM1*, *GSTO1*, *GSTO2*, *GSTP1*, *GSTP2*, *GSTT2*, and *GSTZ1*), indicative of pervasive adaptive evolution. Considering that cytosolic GST genes play critical roles in the detoxification and metabolic activation of xenobiotics (Board and Menon 2013), these changes are likely to reflect previous and ongoing adaptions to diverse environments by mammals. In addition, 31 positively selected sites with radical amino acid changes were identified by gene- and protein-level selection analyses. Interestingly, 73% of the total amino acids under positive selection were found to be located in or close to functional domains. These results indicate that positively selected amino acid changes might play an important role in modulating the specificity or potency of detoxification and antioxidant defenses during mammalian evolution. For example, residue 45 of *GSTA1* is known to be involved in GSH binding (Balogh et al. 2009), while *GSTA1* residues 212 and 215 are close to a residue (216) important for thiolester substrate hydrolysis (Hederos et al. 2004). Residue 107 (a histidine) of *GSTM1* is a second substrate-binding site and has five radical amino acid changes in mammals. Site-directed mutatagenesis of this residue to an aspargine caused a 50% reduction in catalytic activity (Patskovsky et al. 2006).

### Mammalian cytosolic GSTs phylogeny and birth-and-death evolution

The phylogenetic relationships among 333 cytosolic GSTs were reconstructed by two independent methods which gave a similar estimation of mammalian phylogeny. The branch pattern indicated that the omega, theta, and zeta subclasses are ancient in mammals, while the alpha, mu, pi, and sigma subclasses evolved later (Figure 1). This is in agreement with previous studies based on structural and functional data (Armstrong 1997; da Fonseca et al. 2010; Frova 2006). The omega subclass has a cysteine residue at the active site, while the theta and zeta subclasses employ catalytic serine hydroxyl to activate GSH. They are thus predicted to be the progenitors of GSTs (Frova 2006). According to our phylogenetic tree, the omega subclass harbors the most ancient genes (Figure 1). Omega GSTs have strong homology to glutaredoxins, the predicted ancestors of the N-terminal topology of GST (Frova 2006; Oakley 2005). Taken together, it is thus reasonable to assume that the omega subclass evolved earlier. A switch from serine to tyrosine in the alpha, mu, pi, and sigma subclasses is another evolutionary scenario of GSTs which would explain why these four subclasses cluster together in the tree with high bootstrapping (Figure 1). It would appear that the sigma subclass diverged before the mammalian alpha, mu, and pi group due to its presence in invertebrates and vertebrates (Frova 2006). In our study, however, sigma appeared as the sister group of the alpha subclass. This scenario is in line with a previous study (da Fonseca et al. 2010). The mammalian sigma subclass, known as prostaglandin synthases, has a hydrophilic interface with a lock-and-key motif – similar to the alpha, mu, and pi subclasses (Sheehan et al. 2001). Therefore, the observed clustering might be related to their specialized structure and function in mammals. On the other hand, our result also suggests that subclass mu proteins arose most recently. The theta subclass was placed as sister to a clade containing mu, pi, alpha, and sigma subclass with high support – supporting the prediction that alpha, mu, pi, and sigma subclasses arose from the duplication of theta subclass (Armstrong 1997).

The GST gene family has been previously described in various prokaryotes and eukaryotes (Pearson 2005). In this study, we examined cytosolic GST gene repertories in 21 species representing all major mammalian taxa. Our results show an unequal copy number of GST genes across the mammalian phylogeny, as has previously been suggested (da Fonseca et al. 2010; Pearson 2005). For instance, the largest gene expansion was observed in the mouse (30 copies), whereas only 16 GSTs were identified in bowhead whale. This could be due to a lineage or species-specific duplication or deletions of this gene family in mammals. In support of this possibility, a diverse pseudogene proportion was found in mammalian GSTs; ranging from 41% in Yangtze river dolphin and Yangtze finless porpoise to 0% in the African elephant and Florida manatee (Table 1). Studies of gene duplicates have shown that new genes are usually created by gene duplication (Lynch and Conery 2000; Nei and Rooney 2005). Some duplicated genes are maintained in the genome for a long time, while others are nonfunctional or deleted from the genome. This phenomenon is termed the birth-and-death evolution model (Lynch and Conery 2000; Nei and Rooney 2005). We argue that this model is supported by the gene duplication events observed in our data of mammalian GSTs. It is also important to note that extensive duplication (12 paralogous) of pi GSTs was found in the dog, with half being pseudogenes. It has been reported that most duplicated genes tend to experience a brief period of relaxed selection early in their history, often resulting in non-functionalization or pseudogenes (Lynch and Conery 2000). Therefore, the high fraction of pi GSTs pseudogenes in dog further supports a birth-and-death model of GSTs evolution. Notably, seven pi subclass paralogous identified in mouse are apparently intact and functional, as is the case for the naked mole rat mu GSTs (10 copies). This could be the outcome of adaptations to environmental toxins (mouse) and very low oxygen levels in underground burrows (naked mole rat) via diversification of duplicated copies. Moreover, our results reveal extensive positive selection in *GSTP2* along five mammalian lineages (Figure S5), suggestive of episodic selection pressure – possibly in response to changes in xenobiotic exposure. In contrast, there is no evidence of positive selection in *GSTP1* along specific lineages, indicating a functional conservation which is consistent with a critical role of *GSTP1* in ethacrynic acid metabolism in the liver (Henderson et al. 1998). These results reveal that divergent selective regimes occurred in paralogs within a cytosolic GST class.

### Divergent selection of mammalian cytosolic GSTs

Numerous studies have reported that gene family evolution is closely tied with environments. This includes opsin genes in cichlid fish (Henderson et al. 1998), hemoglobins in vertebrates (Nery et al. 2013), keratin-associated proteins in vertebrates (Khan 2014; Sun et al. 2017), and olfactory receptor genes in mammals (Hughes et al. 2018; Niimura et al. 2018). An interesting result of our analysis is the correlation between distinct ecological milieus and cytosolic GST gene copy number (Table 1). For example, in Carnivora 29 genes were identified in dog, while 22 and 25 genes were found in the Weddell seal and Pacific walrus, respectively. In Cetartiodactyla, 21 to 25 genes were found in Artiodactyla, while 16 to 17 genes were found in Cetacea. We, therefore, hypothesize that the dynamic evolution of the cytosolic GST gene family reflects ecological adaptations in aquatic and terrestrial species. We also show that hypoxia-tolerant species with different ecological niches show evidence of divergent selection (Table S6), further suggesting that habitat plays a role in GST gene family evolution.

Marine mammals (cetaceans, pinnipeds, and sirenians) encompass phenotypic convergences that accommodate the challenges of aquatic life. They present a similar respiratory and cardiovascular solutions, such as improved oxygen storage, to low oxygen levels (Ramirez et al. 2007). Our data suggests that their shared adaptations are not reflected in the evolution of the GST gene family. A total of four, three, and one positively selected genes were identified along cetaceans, pinnipeds, and sirenians, respectively (Figure S5). This suggests that there is a difference in the evolutionary history of the GST family in marine mammals. There are several non-mutually exclusive scenarios that could explain this pattern: 1) Antioxidant status is directly related to diving capacity. Marine mammals that perform shallow/short and deep/long divers might experience different oxidative stress challenges, suggesting distinct mechanisms to maintain redox balance (Cantú-Medellín et al. 2011); 2) Pinnipeds and sirenians possess enhanced enzymatic antioxidant capacities, whilst non-enzymatic antioxidant (e.g., levels of glutathione) seems to play an important role in cetaceans, indicating a different strategy for antioxidant defenses adaptation (Ninfali and Aluigi 1998; Wilhelm Filho et al. 2002); 3) We do not rule out the potential impact of different number of species and the length of branches used in this study. As an ever-increasing amount of high-quality genome assemblies are generated, future studies is likely to resolve these issues.

### Oxidative stress adaptation in cetaceans

Cetaceans are faced by chronic oxidant stress stemming from chemical pollutants in aquatic environment and reoxygenation following hypoxia (diving) (Valavanidis et al. 2006; Li and Jackson 2002). Therefore, it might be expected that cetaceans have a large GST repertoire. However, we found that the number of cytosolic GST genes in cetaceans was significantly smaller than terrestrials (Figure 3C). The contraction of alpha GSTs is striking, with only two functional *GSTAs* (*GSTA1* and *GSTA4*) in cetaceans. Similarly, five mu GSTs were identified in cetaceans, but only one gene (*GSTM1*) appears intact compared with 5-10 GSTM genes in other mammals (Figure 2). We observed that ancestral branches of cetaceans also showed reduced GST repertoires, which suggested that contraction of cetacean GSTs could be related to aquatic adaptations after the divergence of cetacea from artiodactyla approximately 55 Mya (Thewissen et al. 2007). The presence of four *GSTM* pseudogenes provides further evidence for the contraction of the cetacean locus from a large GST family in the ancestral cetacean genome. Moreover, these gene losses in cetaceans probably a consequence of relaxed selection after adaptations of aquatic environment (Table S7). This raises the probability of an alternative gene family responsible for enhanced oxidative stress resistance or perhaps selection on particular genes responsible for detecting toxicants and activating oxidative defenses, rather than a large gene repertoire. Regarding the former explanation, it has been reported that the peroxiredoxin (*PRDX*) and glutathione peroxidase (*GPX*) gene families have expanded in whale lineages (Yim et al. 2014; Zhou et al. 2018). On the other hand, it is also possible that the retained GSTs in cetaceans have improved or essential antioxidative properties. For example, single nucleotide polymorphisms that cause amino acid substitutions in *GSTA1* alters its activity towards xenobiotics (Coles and Kadlubar 2005). *GSTA4* protects against oxidative stress by clearing toxic lipid peroxidation by-products (Hayes et al. 2005). *GSTM1* is critical for the detoxification of various oxidants, as well as carcinogens and toxins (Mcilwain and Townsend DMTew 2006). Supporting an essential role of these cytosolic GST genes (*GSTA1*, *GSTA4*, and *GSTM1*), human and rodent studies have reported that these GSTs are also present in mitochondria where they likely play a role in protecting against mitochondrial injury during oxidative stress (Gallagher et al. 2006; Raza et al. 2002) We speculate that the cetacean homologs, in particular *GSTA1* which was intact in all seven cetaceans, play an essential role in protecting against oxidative stress by localizing to both the cytosol and mitochondria. That is, complete loss of these genes are not tolerated. We also show evidence of positive selection on the retained GSTs in cetaceans. These observations lead us to speculate that widely dispersed xenobiotics in aquatic ecosystems and high oxidative stress drive the adaptive evolution of retained functional and essential GST genes in cetaceans.

Previous studies demonstrated that gene loss in cetaceans could play an important role for natural phenotypic adaptations. For example, loss of genes with hair- and epidermis-related functions contribute to their unique skin morphology, a thicker epidermis and hairlessness (Sharma et al. 2018). Some investigators have reported that gene loss may carry detrimental fitness consequences in modern environments. An intriguing recent example includes the loss of paraoxonase 1 (*PON1*) in marine mammals which likely eliminates their main defense against neurotoxicity from man-made organophosphorus compounds (Meyer et al. 2018). Loss of GST genes in cetaceans may also have negative consequences. For instance, *Gsta3* knockout mice are not only sensitive to acute cytotoxic and genotoxic effect of aflatoxin B1 (Zoran et al. 2010), but also have increased oxidative stress marker levels (Crawford et al. 2017), suggesting that *GSTA3* loss in cetaceans might also weaken their oxidative damage defenses. Nevertheless, more research is necessary to validate whether a cetacean gene loss event is deleterious.

In conclusion, we here characterized the cytosolic glutathione-*S*-transferase (GST) gene family in 21 mammalian species. In particular, our study shows that the gene family has contracted in cetaceans despite the important role of GST genes in the protection against various stressors. Our findings add another piece to the puzzle of understanding how ceteaceans adapt to an oxygen-poor aquatic environments. An ever-increasing amount of genomic research data and associated tools is likely to address this important research question.

## ACKNOWLEDGEMENTS

This work was financially supported by the National Key Program of Research and Development, Ministry of Science and Technology of China (grant no. 2016YFC0503200 to G.Y. and S.X.), the Key Project of the National Natural Science Foundation of China (NSFC) (Grant no. 31630071 to G.Y.), the National Natural Science Foundation of China (NSFC) (grant nos. 31570379, 31772448 to S.X., grant no. 31872219 to W.R.), the Priority Academic Program Development of Jiangsu Higher Education Institutions (PAPD), and the China Postdoctoral Science Foundation (grant no. 2018M642278 to R.T.).

---

**Figure S1.**
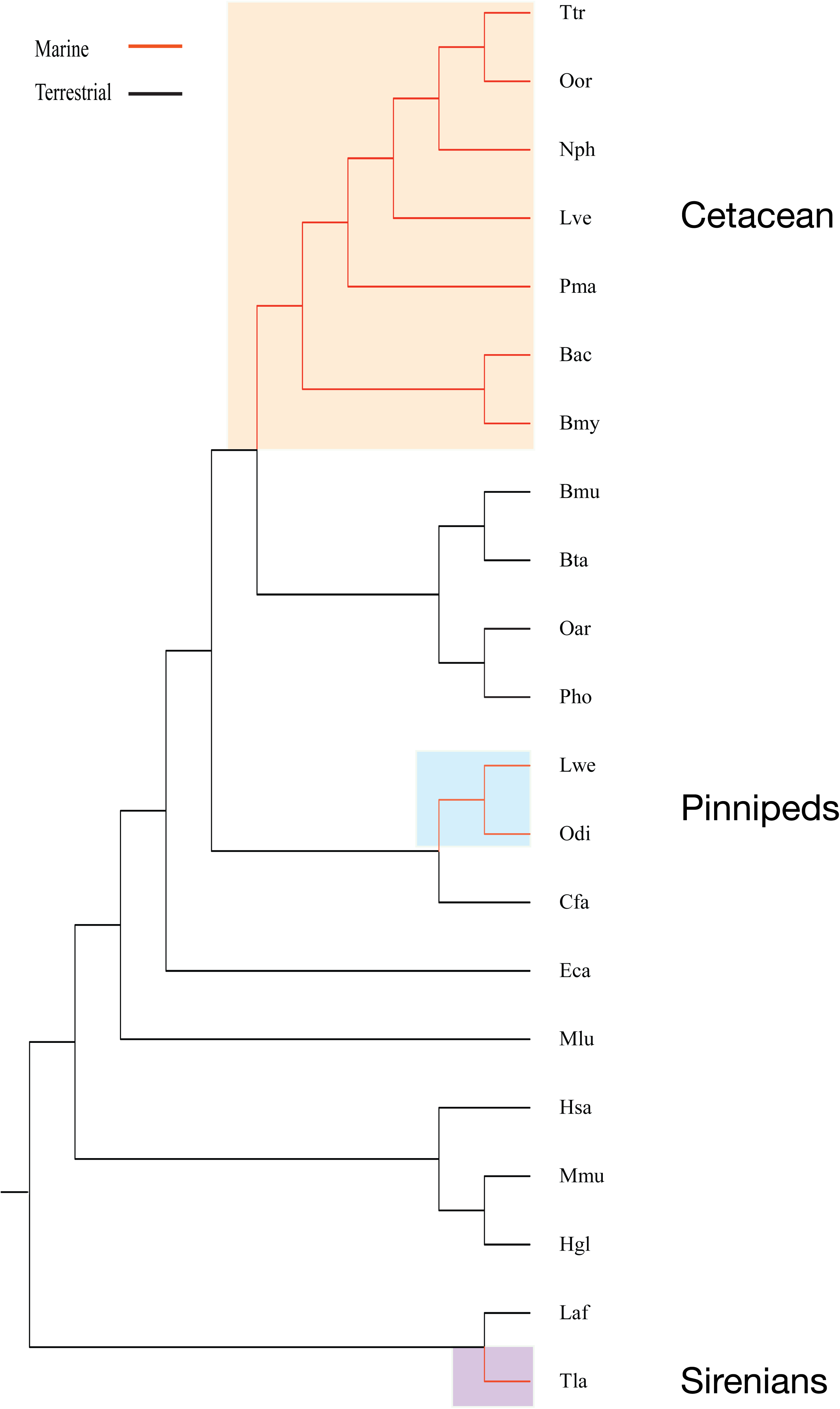
Mammalian species phylogeny (Douzery et al. 2014) indicating each of the clade partitions tested by the branch-site and clade models.

**Figure S2.**
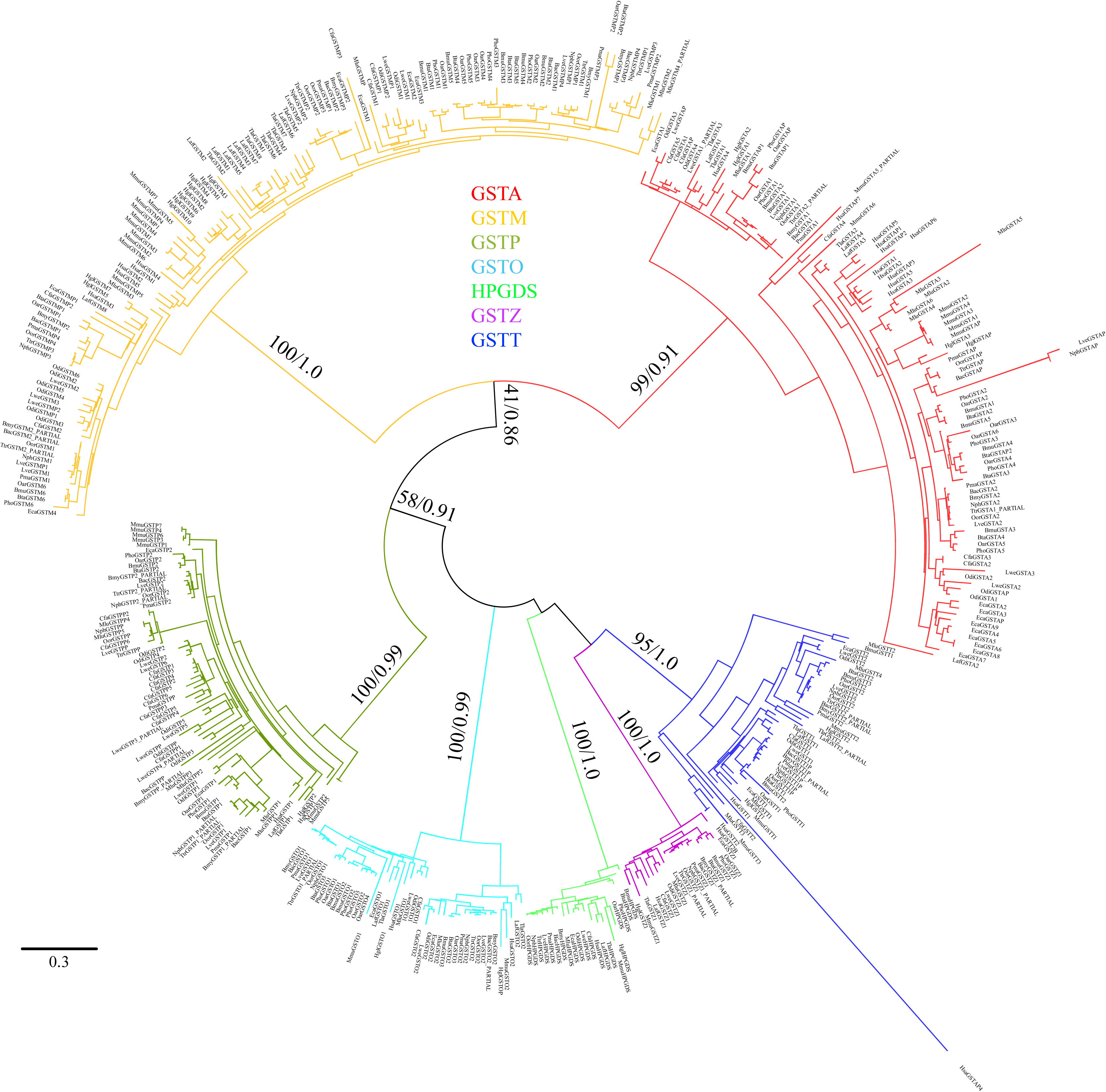
Phylogenetic tree of the GST gene family in mammals. Shown is the maximum likelihood and Bayesian tree built using RaxML and Mrbayes methods with the multiple alignments of 448 GST nucleotide sequences from 21 mammalian species. The bootstrap values and posterior probability are shown at nodes. The clades of alpha, mu, pi, omega, sigma, zeta, theta are shown in red, orange, turquoise, azure, green, purple, and blue, respectively.

**Figure S3.**
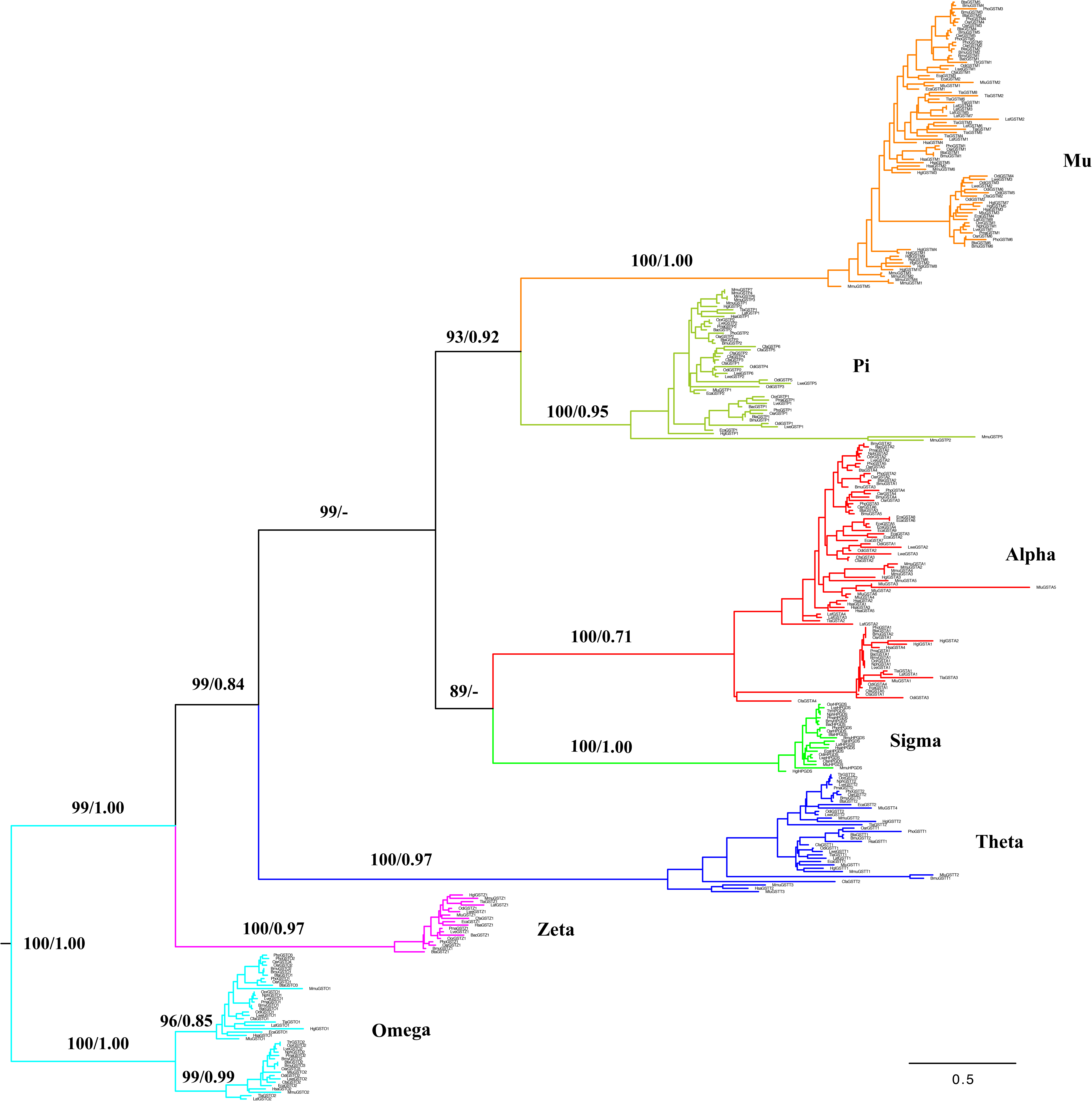
Phylogenetic tree of GST gene family based on 333 mammalian protein sequences. Numbers of the nodes correspond to maximum likelihood bootstrap support values and Bayesian posterior probabilities. The clades of alpha, mu, pi, omega, sigma, zeta, theta are shown in red, orange, turquoise, azure, green, purple, and blue, respectively.

**Figure S4.**
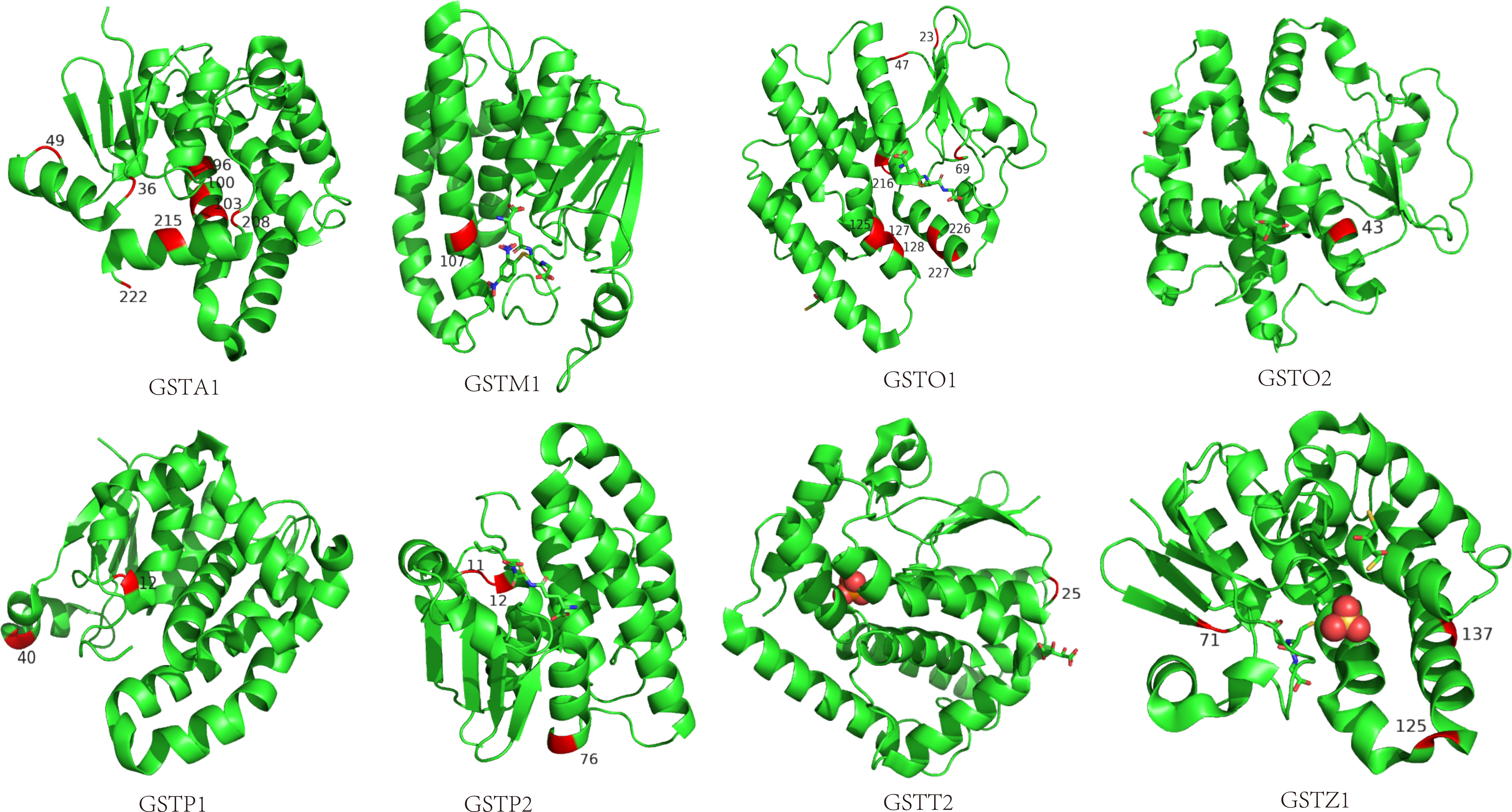
Positively selected sites are shown in crystal structure with red. The crystal structures of GSTA1 (1gsd), GSTM1 (1XW6), GSTO1 (5V3Q), GSTO2 (3Q18), GSTP1 (5X79), GSTP2 (P46425), GSTT2 (4MPF), GSTZ1 (2CZ2) were taken from the Protein Data Bank (http://www.rcsb.org/pdb).

**Figure S5.**
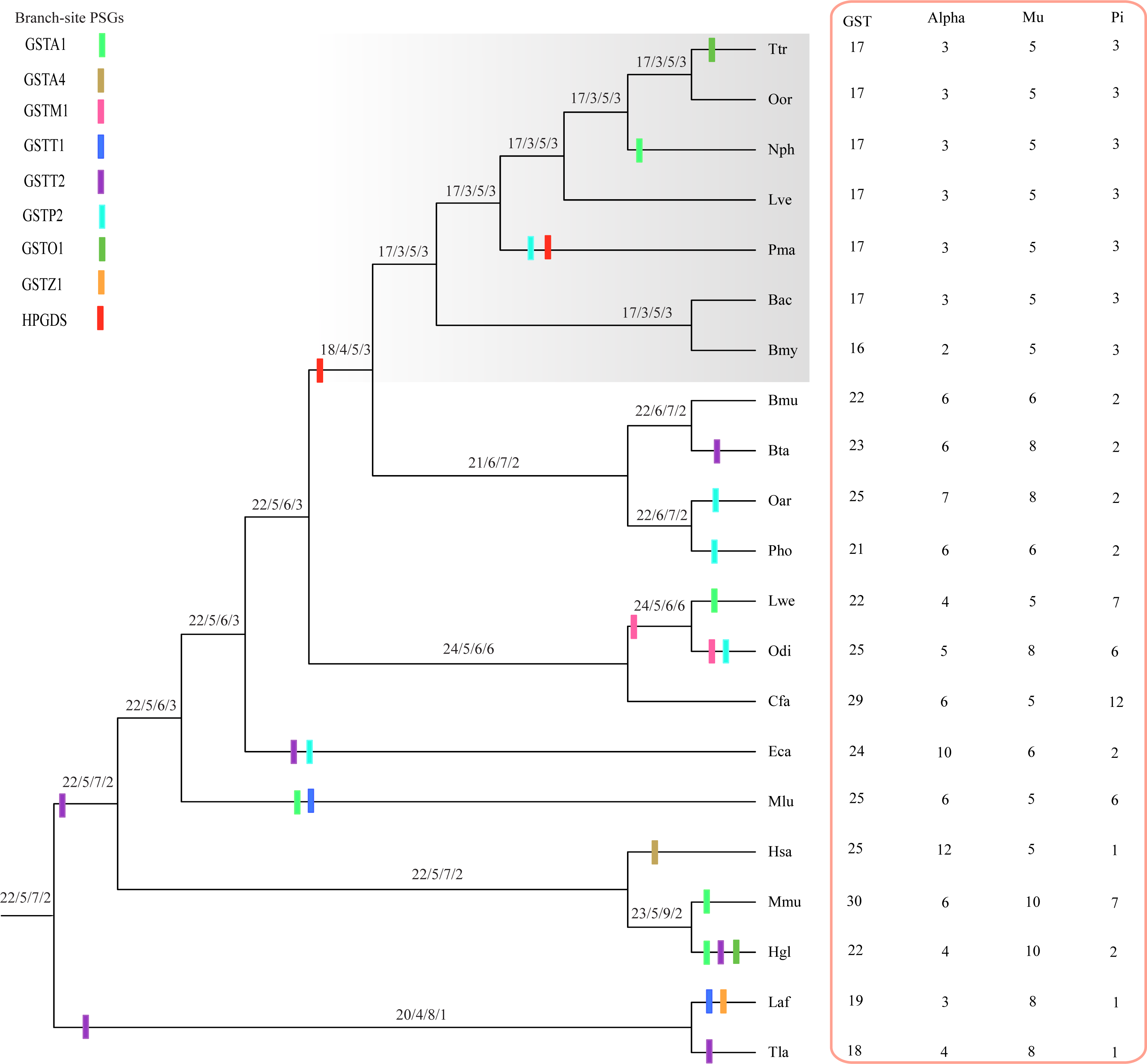
Evidence for lineage-specific positive selection in mammalian GSTs. Inset shows the gene number of GSTs, alpha, mu and pi class inferred to have occurred gain and loss along mammalian lineages.

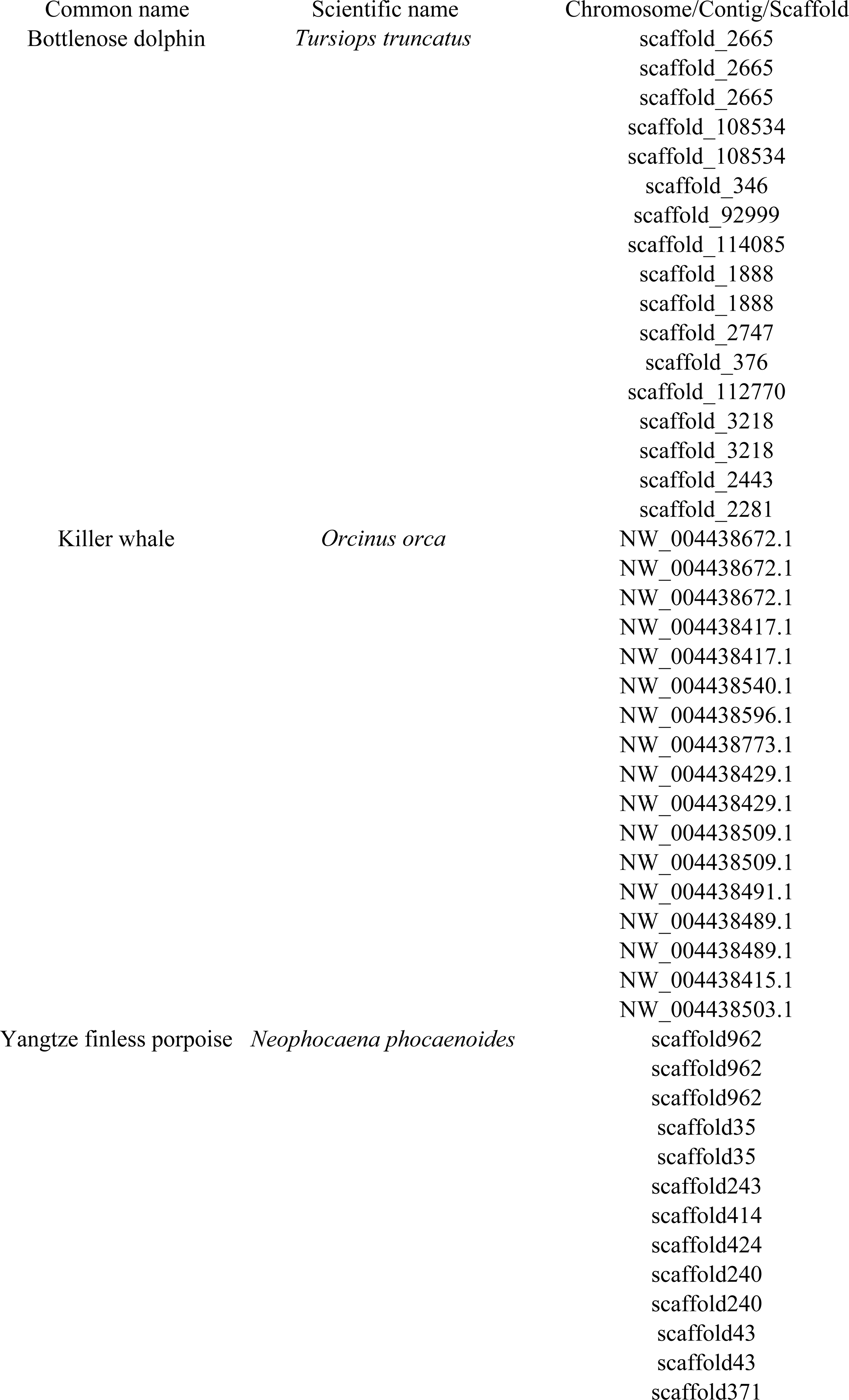
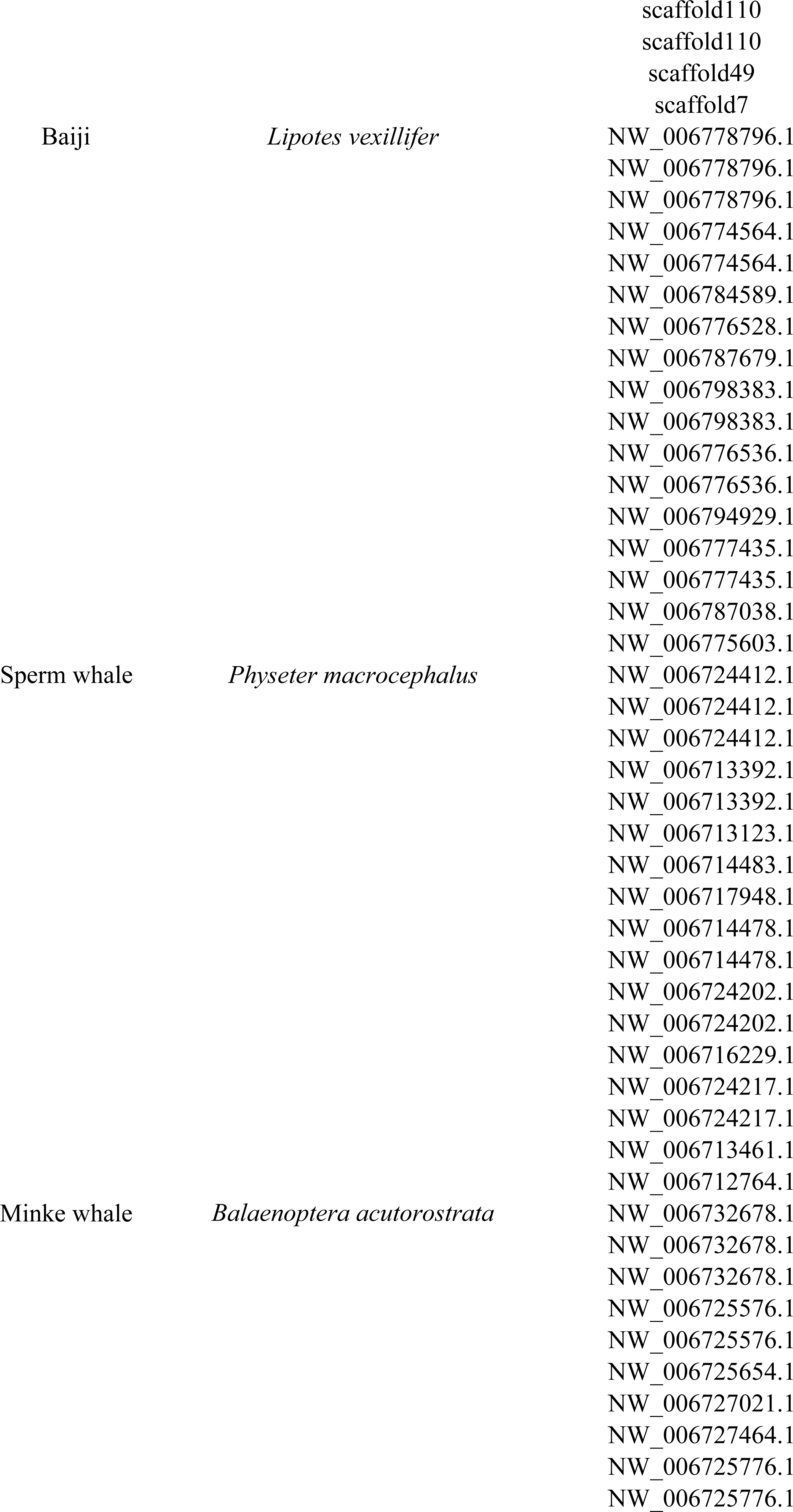
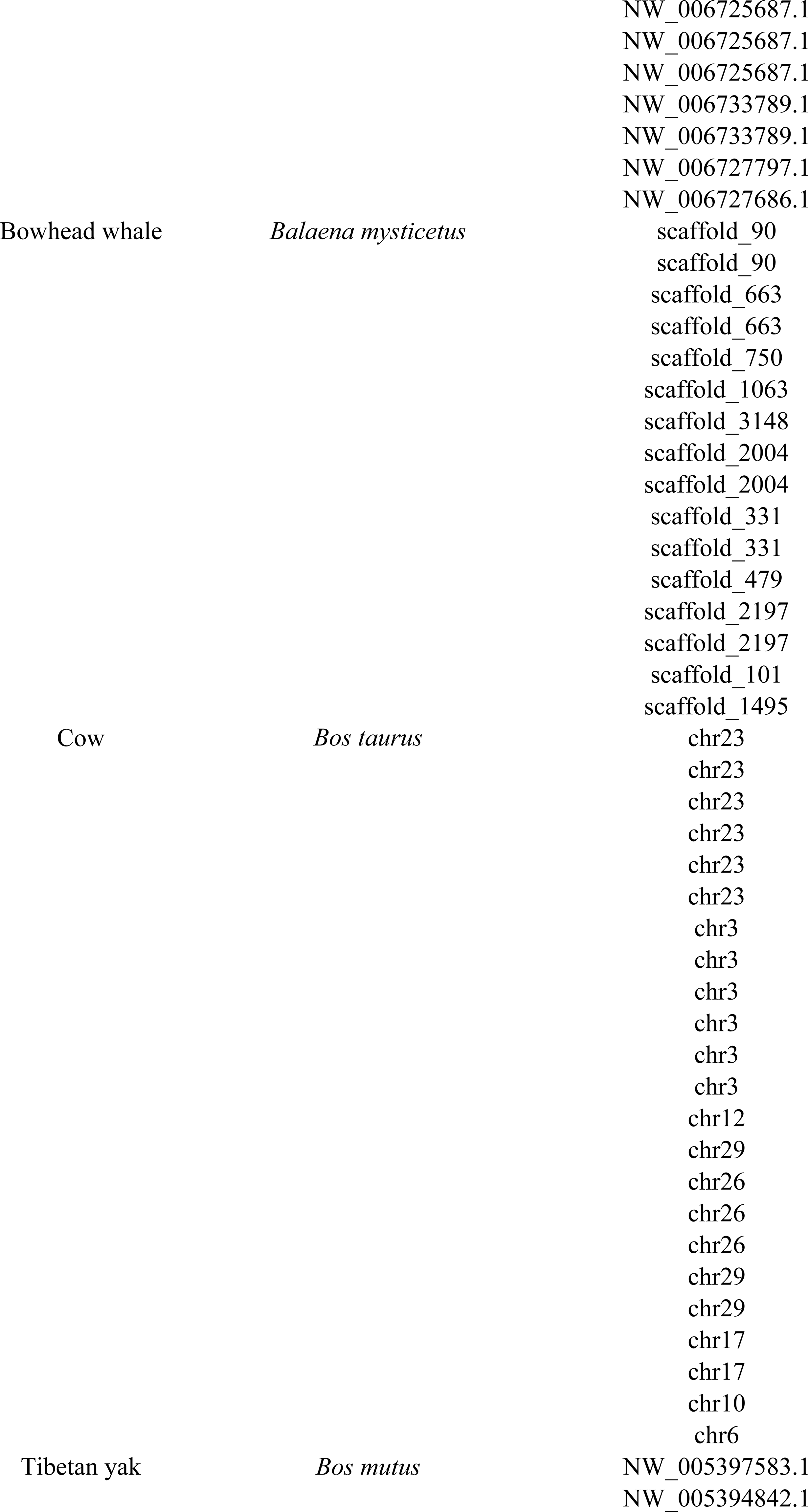
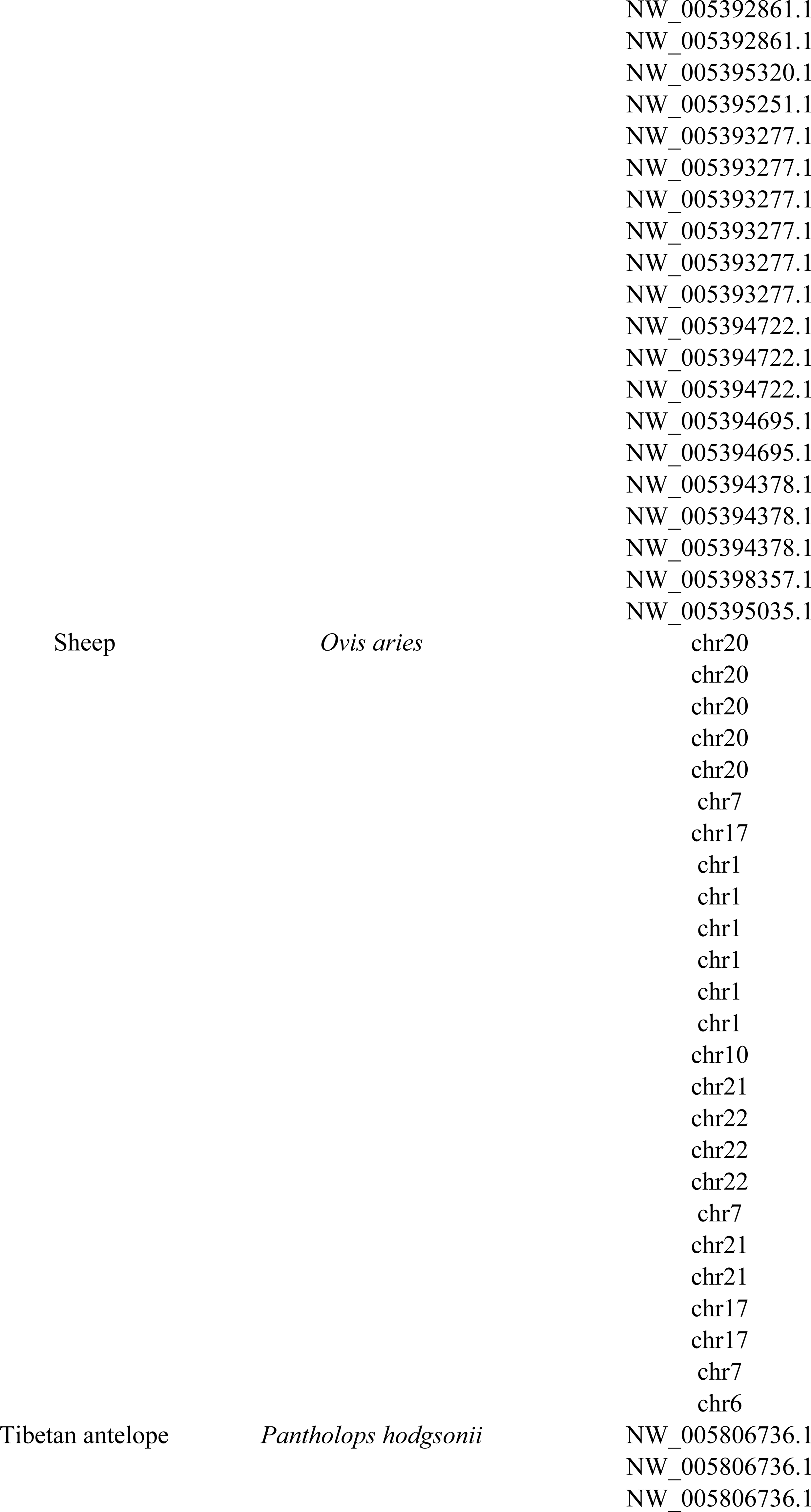
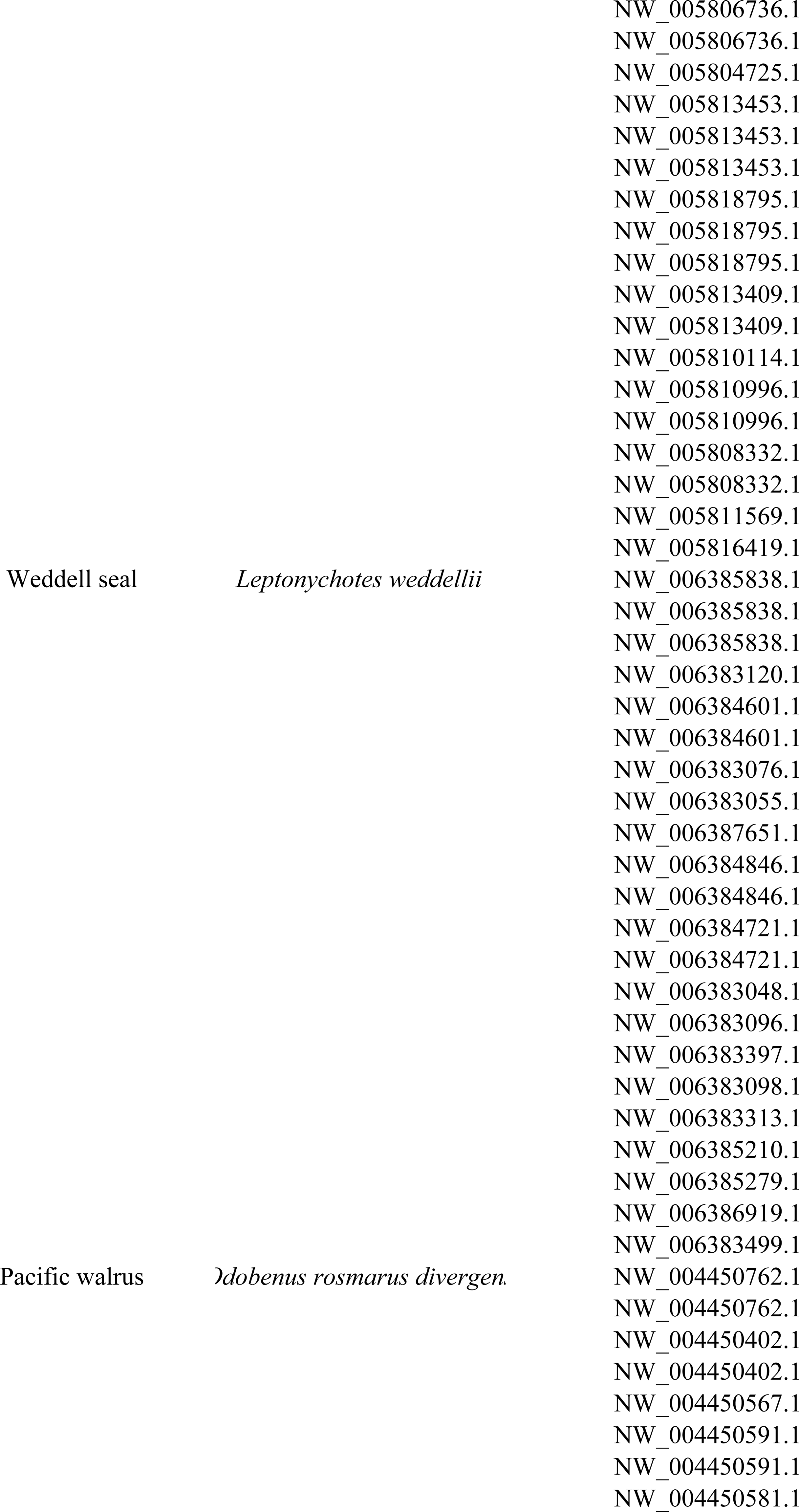
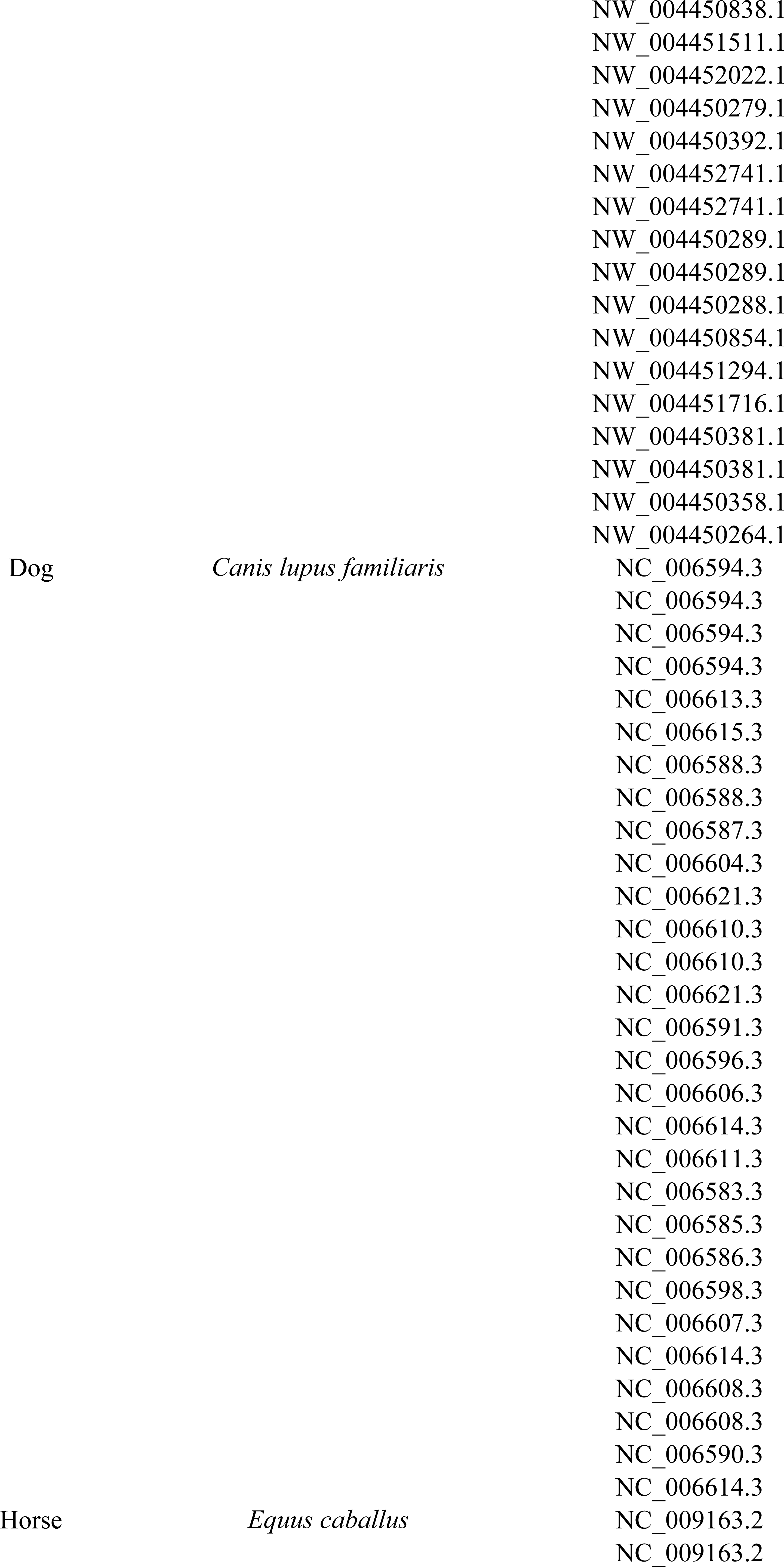
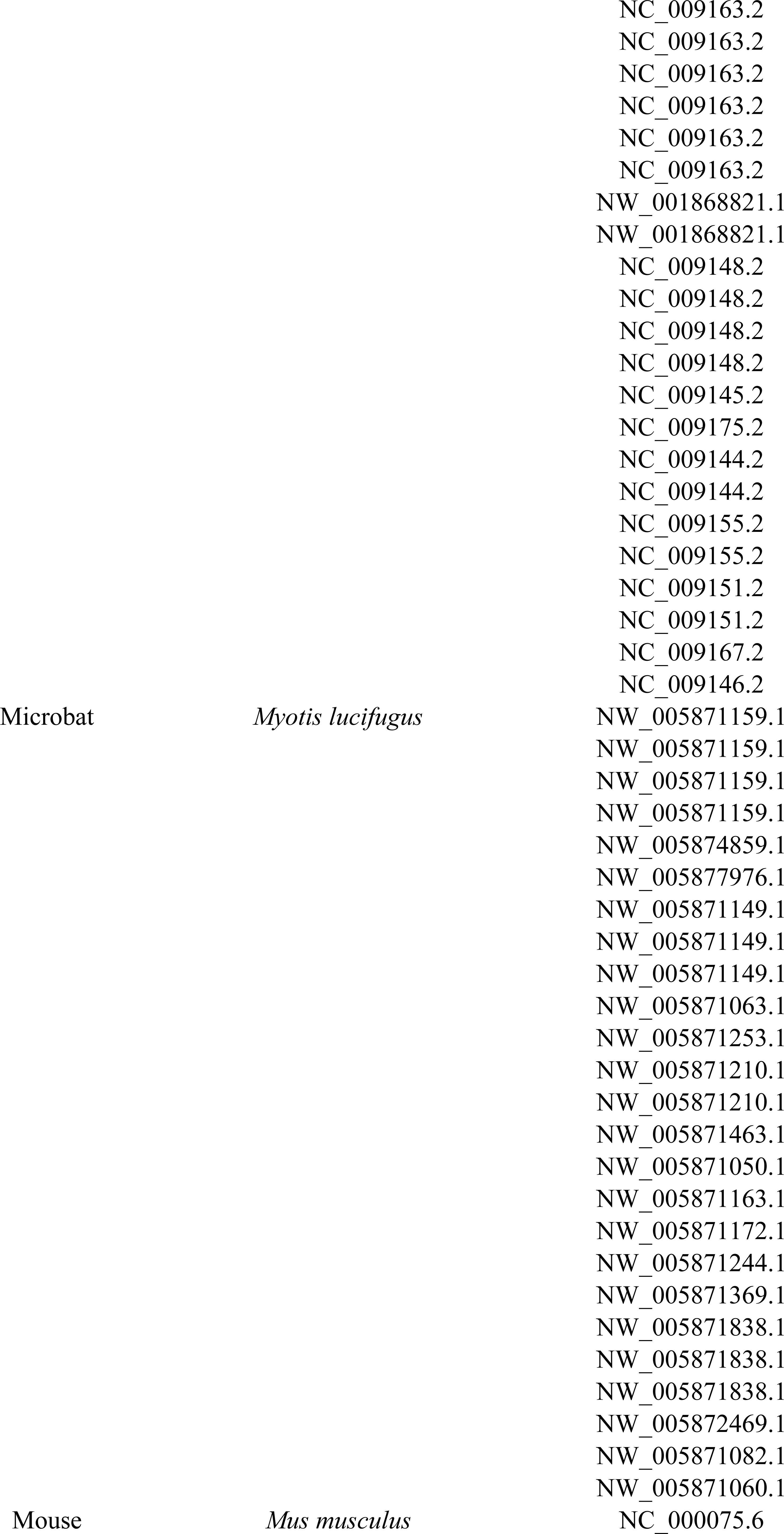
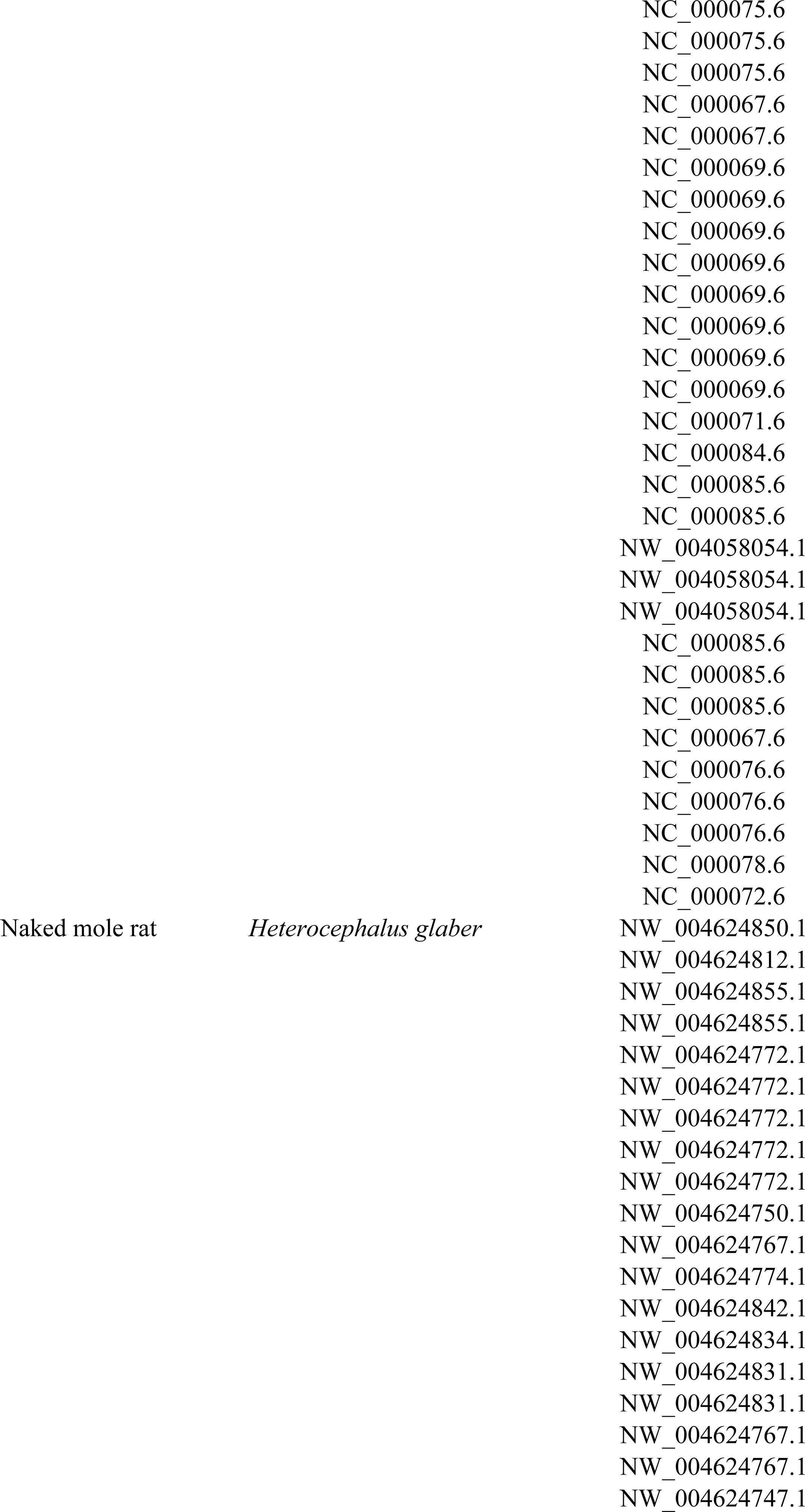
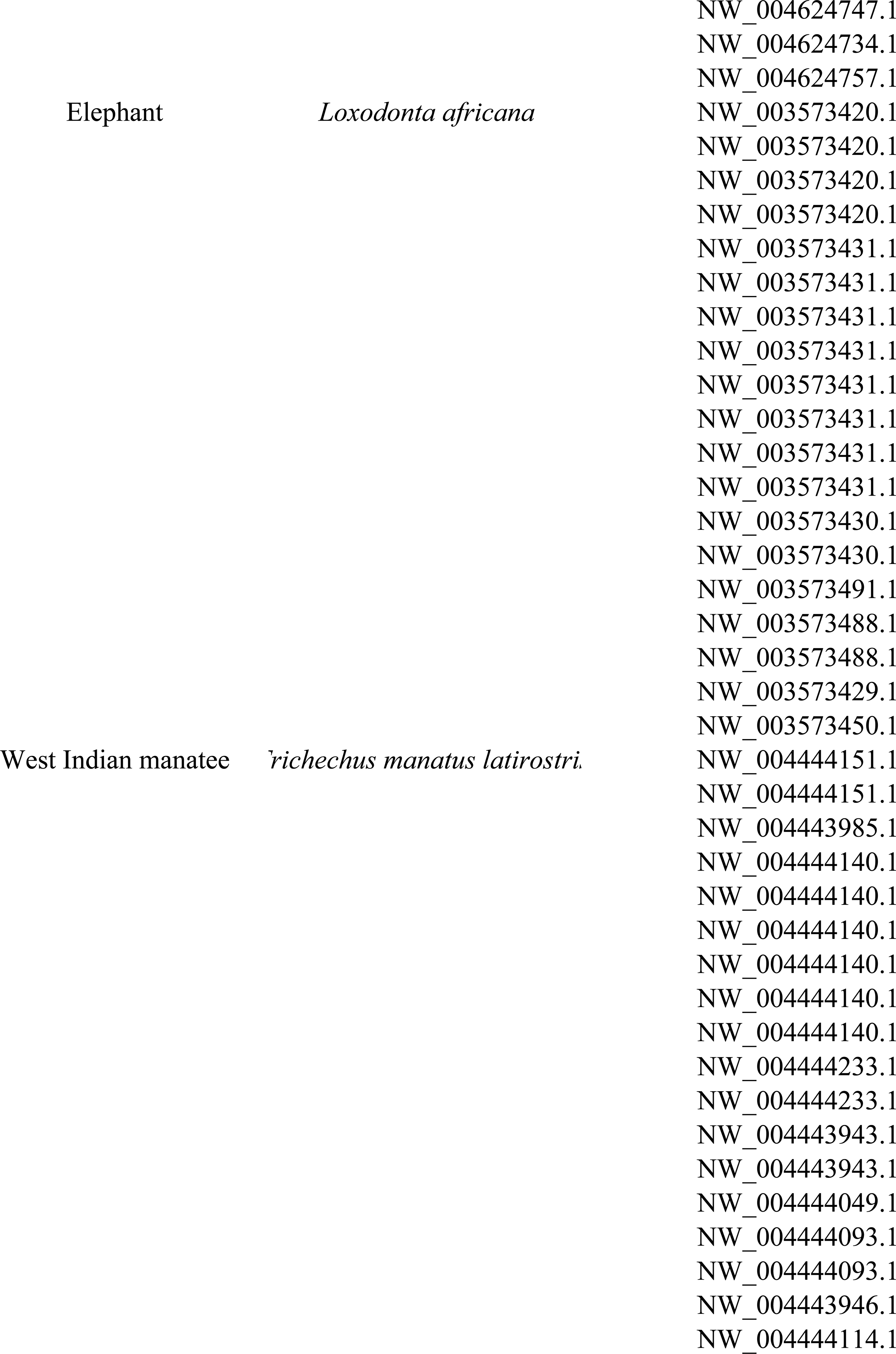
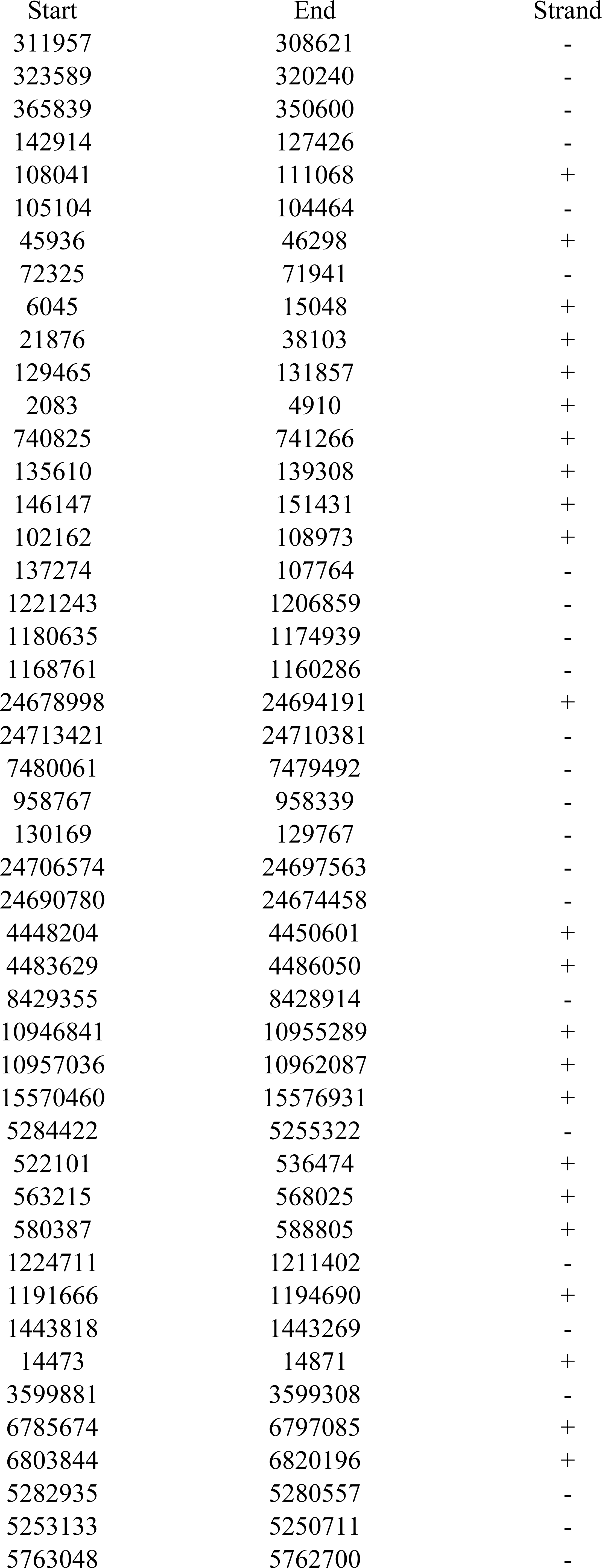
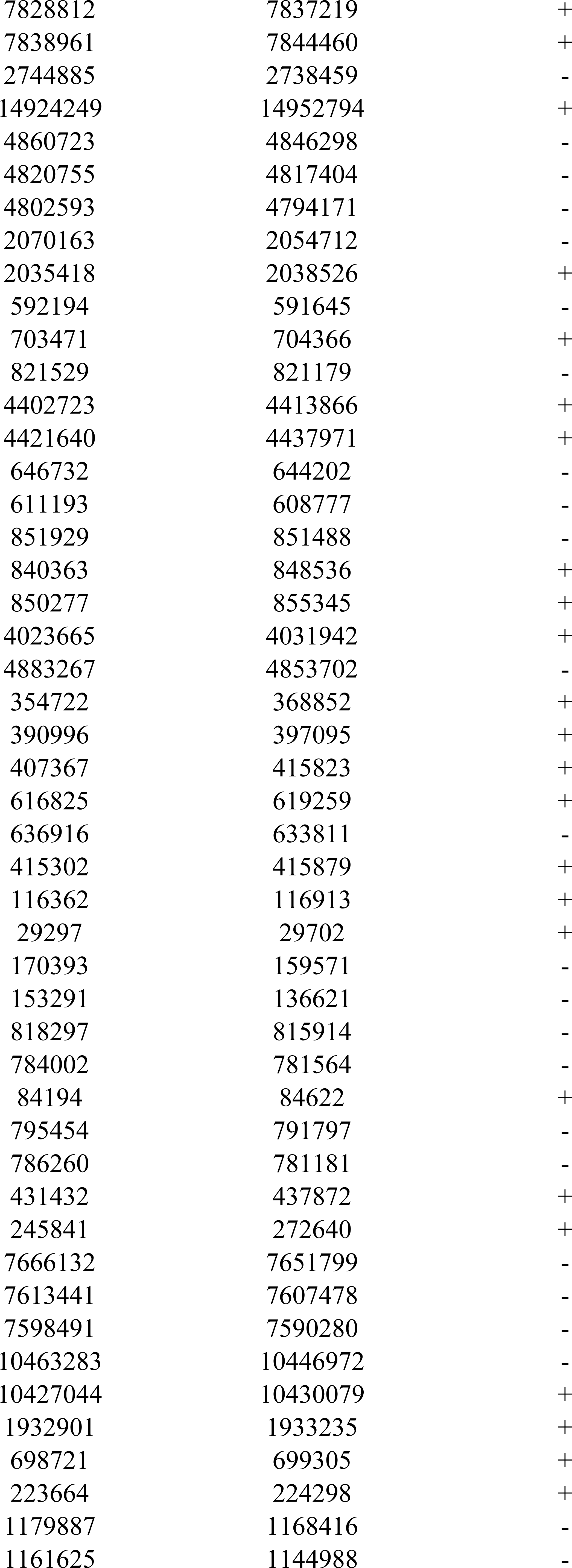
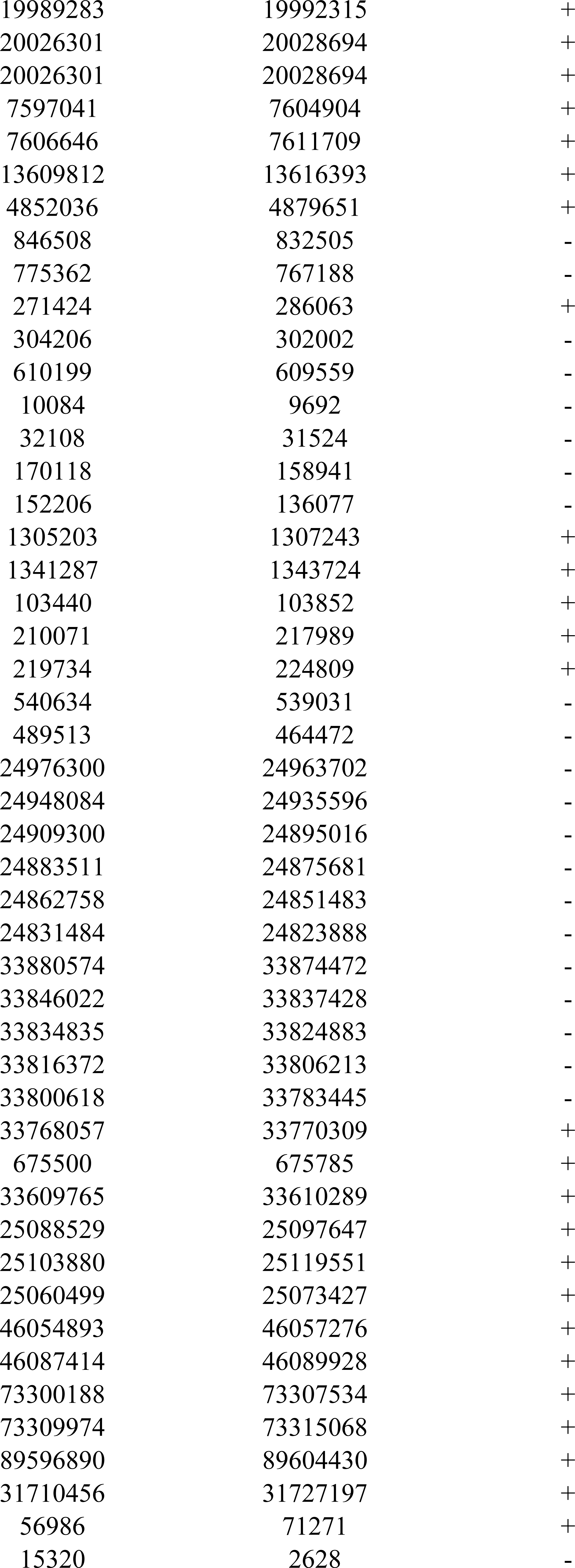
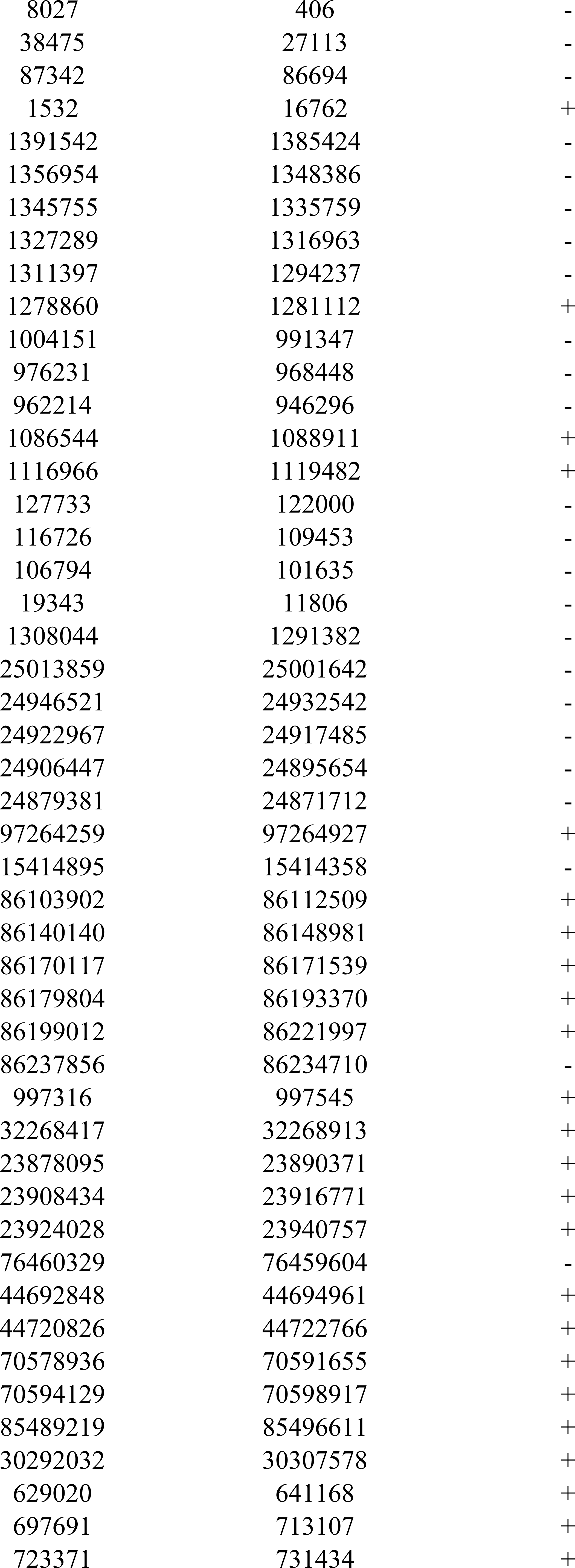
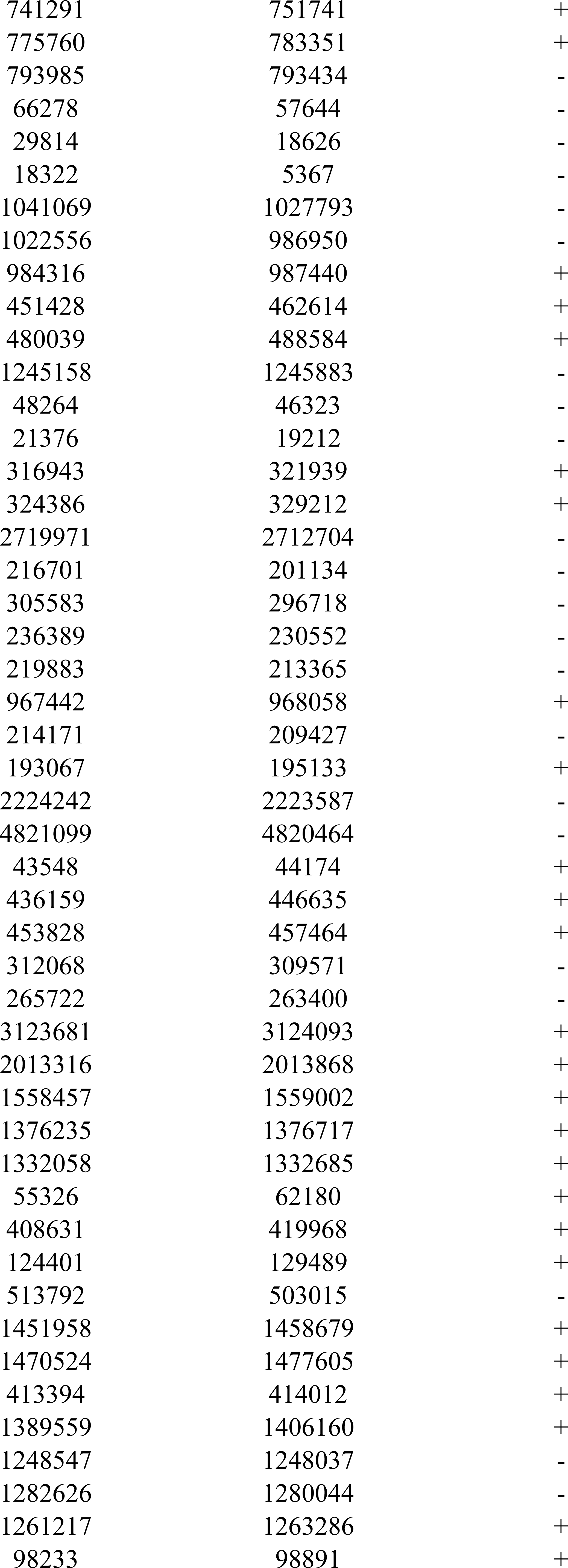
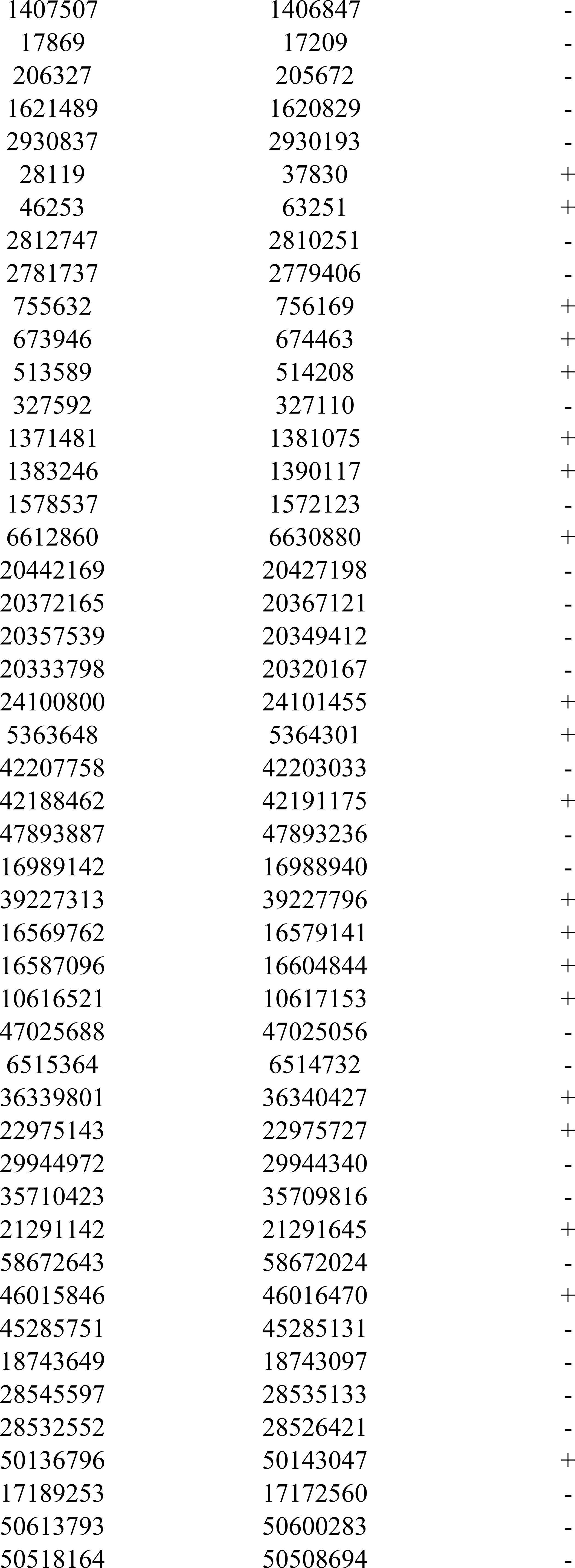
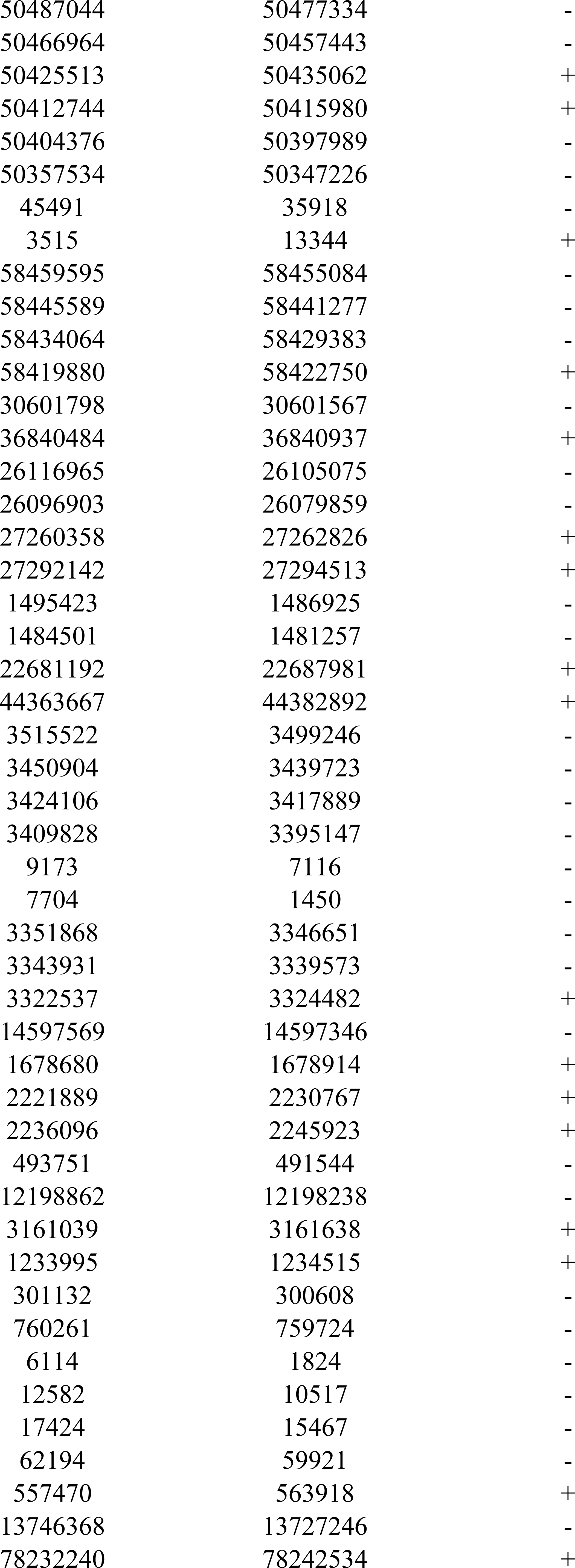
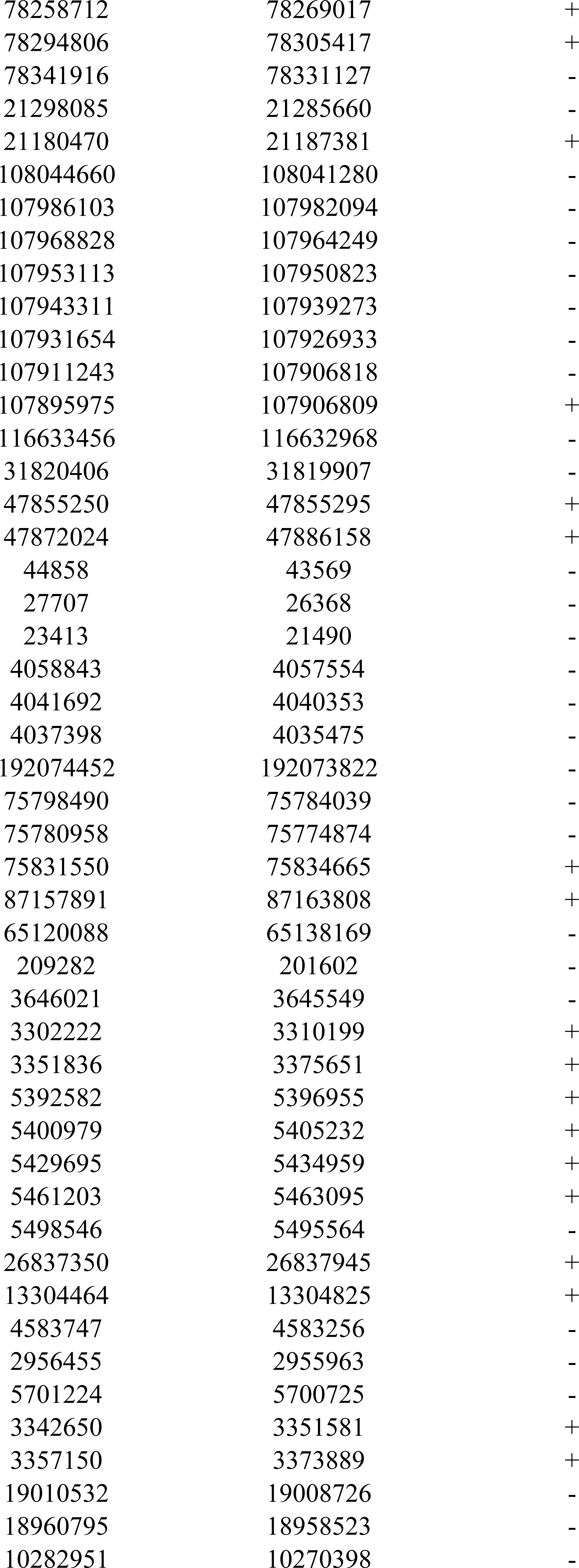
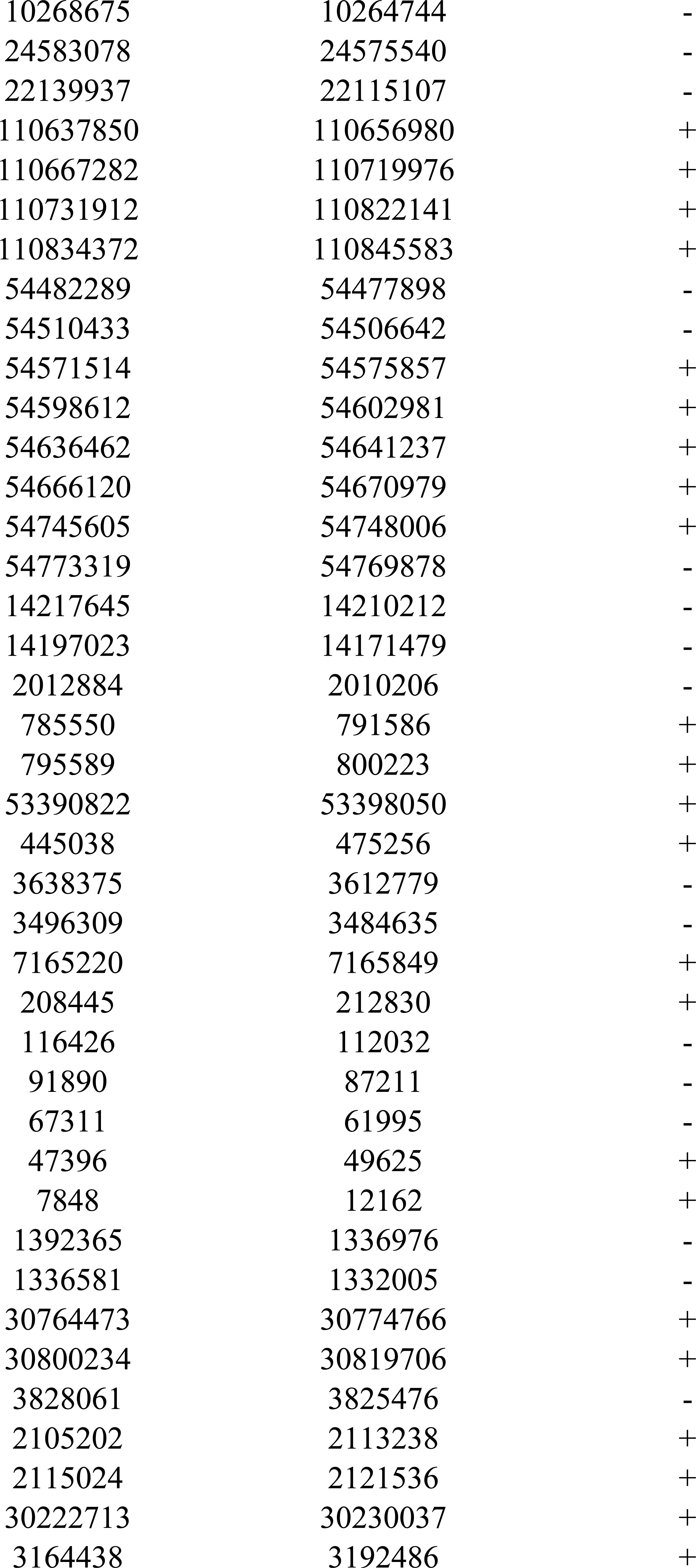
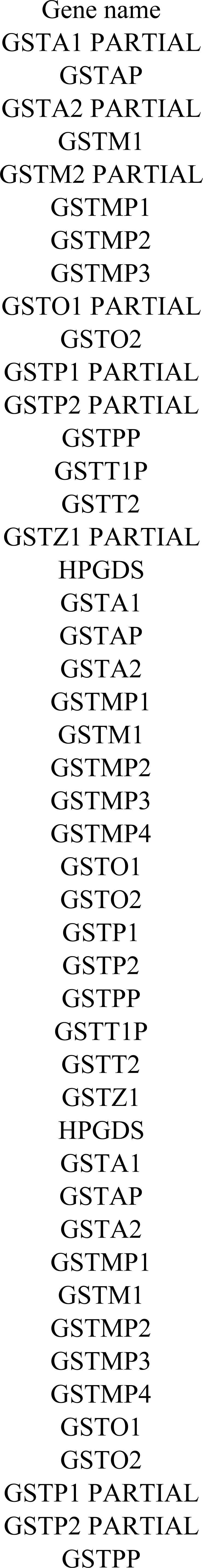
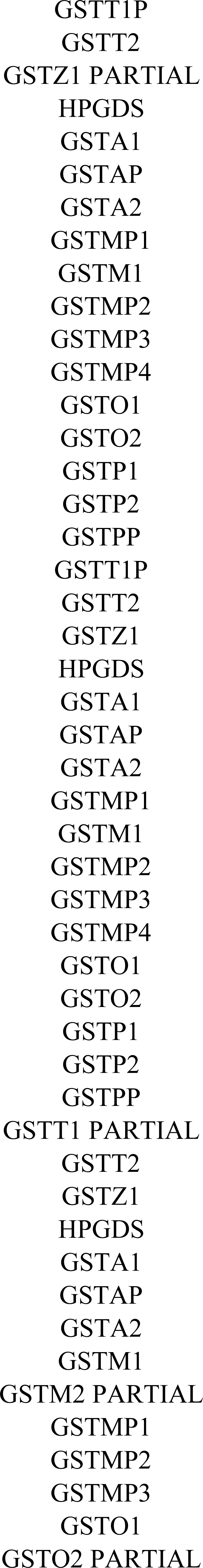
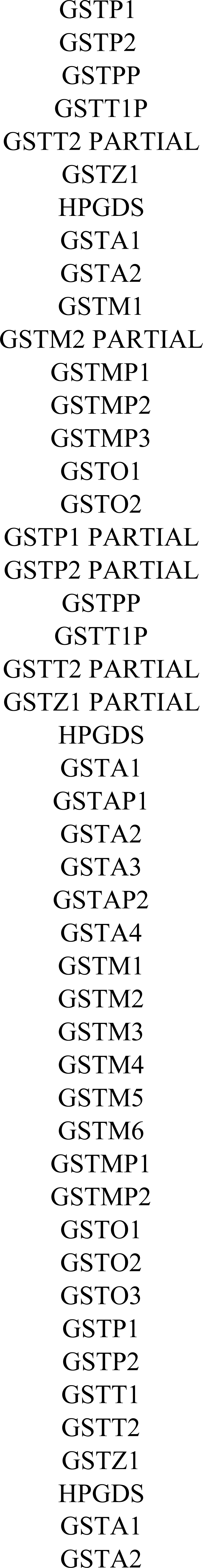
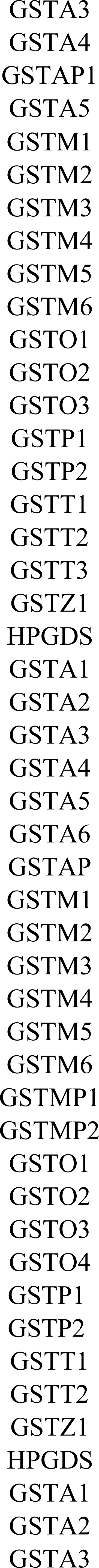
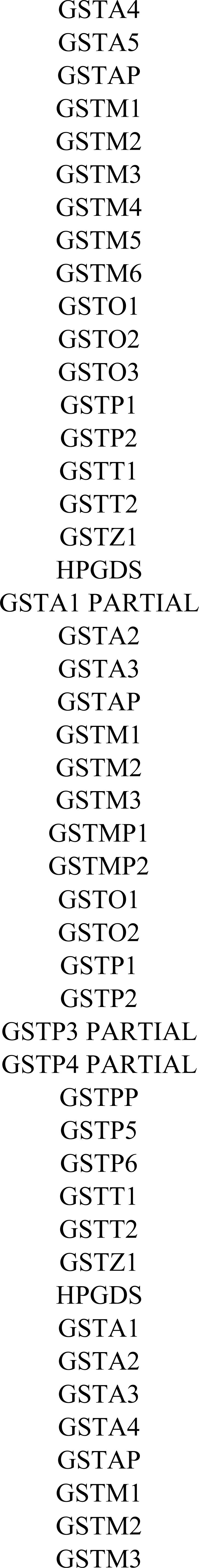
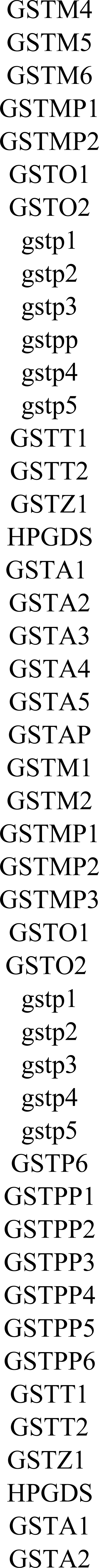
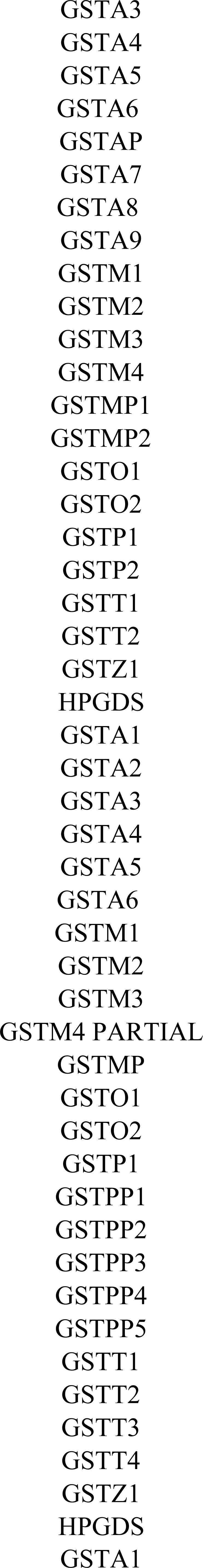
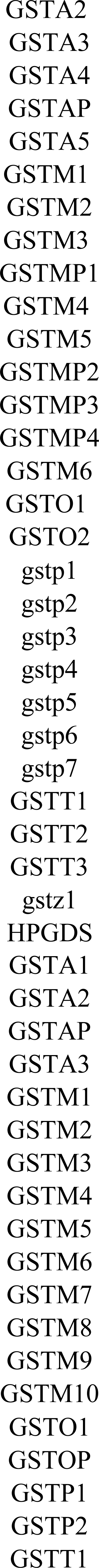
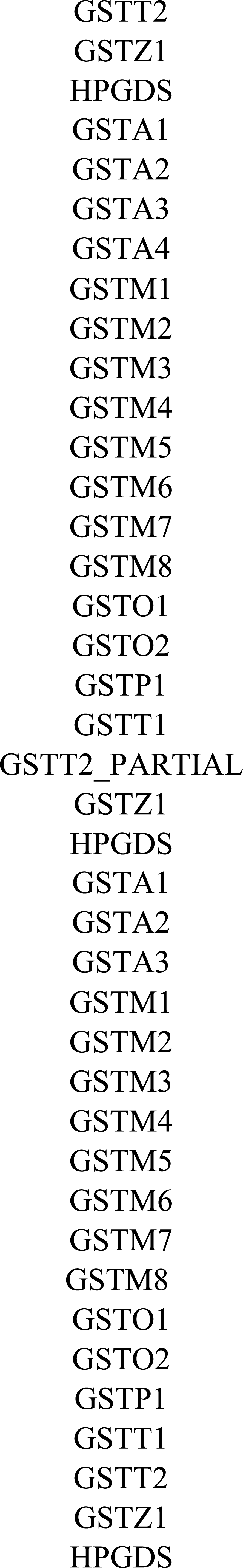

**Table S2.** Genomic information of species used in this study.

| Common name | Latin name | Sequencing technology | Sequence coverage (×) | Scaffold N50 | NCBI Genomes version | RefSeq Assembly Accession | Genome resource | Assembly Completeness in BUSCO (%) |
| --- | --- | --- | --- | --- | --- | --- | --- | --- |
| **Bottlenose dolphin** | *Tursiops truncatus* | Sanger, 454 FLX, Illumina HiSeq | 30 | 116,287 | Ttru_1.4 | GCF_001922835.1 | NCBI | 92.9 |
| **Killer whale** | *Orcinus orca* | Illumina HiSeq | 200 | 12,735,091 | Oorc_1.1 | GCF_000331955.2 | NCBI | 99.3 |
| **Yangtze river dolphin** | *Lipotes vexillifer* | Illumina HiSeq | 115 | 2,419,148 | Lipotes_vexillifer_v1 | GCF_000442215.1 | NCBI | 98.8 |
| **Yangtze finless porpoise** | *Neophocaena asiaeorientalis* | Illumina HiSeq | 106 | 6,341,296 | Neophocaena_asiaeorientalis_v1 | GCF_003031525.1 | NCBI | 99.1 |
| **Sperm whale** | *Physeter macrocephalus* | Illumina HiSeq | 75 | 427,290 | Physeter_macrocephalus_2.0.2 | GCF_000472045.1 | NCBI | 99.5 |
| **Minke whale** | *Balaena acutorostrata* | Illumina HiSeq | 92 | 12,843,668 | BalAcu1.0 | GCF_000493695.1 | NCBI | 99.1 |
| **Bowhead whale** | *Balaena mysticetus* | Illumina HiSeq | 154 | 877,000 | - | - | http://www.bowhead-whale.org | 74.6 |
| **Cow** | *Bos taurus* | Sanger, PacBio RS II | 19 | 6,806,220 | Btau_5.0.1 | GCA_000003205.6 | NCBI | 99.5 |
| **Tibetan yak** | *Bos mutus* | Illumina HiSeq, | 130 | 1,407,960 | BosGru_v2.0 | GCF_000298355.1 | NCBI | 98.1 |
| **Sheep** | *Ovis aries* | Illumina GA<br>Illumina<br>GAII, 454,<br>PacBio RSII | 166 | 100,009,711 | Oar_v4.0 | GCF_000298735.2 | NCBI | 99.7 |
| **Tibetan antelope** | *Pantholops hodgsonii* | Illumina GA | 67 | 2,772,860 | PHO1.0 | GCF_000400835.1 | NCBI | 98.6 |
| **Dog** | *Canis lupus familiaris* | Sanger | 7 | 45,876,610 | CanFam3.1 | GCF_000002285.3 | NCBI | 99.2 |
| **Weddell seal** | *Leptonychotes weddellii* | Illumina<br>HiSeq | 82 | 904,031 | LepWed1.0 | GCF_000349705.1 | NCBI | 87.3 |
| **Pacific walrus** | *Odobenus rosmarus divergens* | Illumina<br>HiSeq | 200 | 2,616,778 | Oros_1.0 | GCF_000321225.1 | NCBI | 99.6 |
| **Horse** | *Equus caballus* | Sanger | 6.8 | 46,749,900 | EquCab2.0 | GCF_000002305.2 | NCBI | 99.7 |
| **Microbat** | *Myotis lucifugus* | Sanger | 7 | 4,293,315 | Myoluc2.0 | GCF_000147115.1 | NCBI | 97.4 |
| **Mouse** | *Mus musculus* | Sanger | - | 52,589,046 | GRCm38.p4 | GCF_000001635.24 | NCBI | 99.9 |
| **Naked mole rat** | *Heterocephalus glaber* | Illumina<br>HiSeq | 90 | 20,532,749 | HetGla_female_1.0 | GCF_000247695.1 | NCBI | 99.3 |
| **African bush elephant** | *Loxodonta africana* | Sanger, ABI | 7 | 46,401,353 | Loxafr3.0 | GCF_000001905.1 | NCBI | 98.7 |
| **Florida manatee** | *Trichechus manatus latirostris* | Illumina<br>HiSeq | 150 | 14,442,683 | TriManLat1.0 | GCF_000243295.1 | NCBI | 98.1 |

**Table S3.** Amino acid sequence similarity (identity) between and within cytosolic GST subclass in seven cetaceans.

| **HPGDS** |  |  |  |  |  |  |  |  |  |  |
| --- | --- | --- | --- | --- | --- | --- | --- | --- | --- | --- |
| **GSTA1** | 25.0% |  |  |  |  |  |  |  |  |  |
| **GSTA2** | 21.4% | 56.0% |  |  |  |  |  |  |  |  |
| **GSTM1** | 25.5% | 25.2% | 20.0% |  |  |  |  |  |  |  |
| **GSTO1** | 17.2% | 19.1% | 17.2% | 15.7% |  |  |  |  |  |  |
| **GSTO2** | 17.6% | 16.6% | 15.6% | 15.3% | 62.3% |  |  |  |  |  |
| **GSTP1** | 24.5% | 30.7% | 27.2% | 31.1% | 15.7% | 20.8% |  |  |  |  |
| **GSTP2** | 23.8% | 30.9% | 27.9% | 30.8% | 18.9% | 21.2% | 82.0% |  |  |  |
| **GSTT1** | 15.4% | 21.1% | 17.8% | 18.6% | 16.6% | 18.9% | 21.5% | 22.3% |  |  |
| **GSTZ1** | 19.9% | 21.3% | 21.3% | 15.6% | 21.2% | 21.0% | 21.1% | 21.0% | 25.5% |  |
|  | HPGDS | GSTA1 | GSTA2 | GSTM1 | GSTO1 | GSTO2 | GSTP1 | GSTP2 | GSTT1 | GSTZ1 |

**Table S4.**
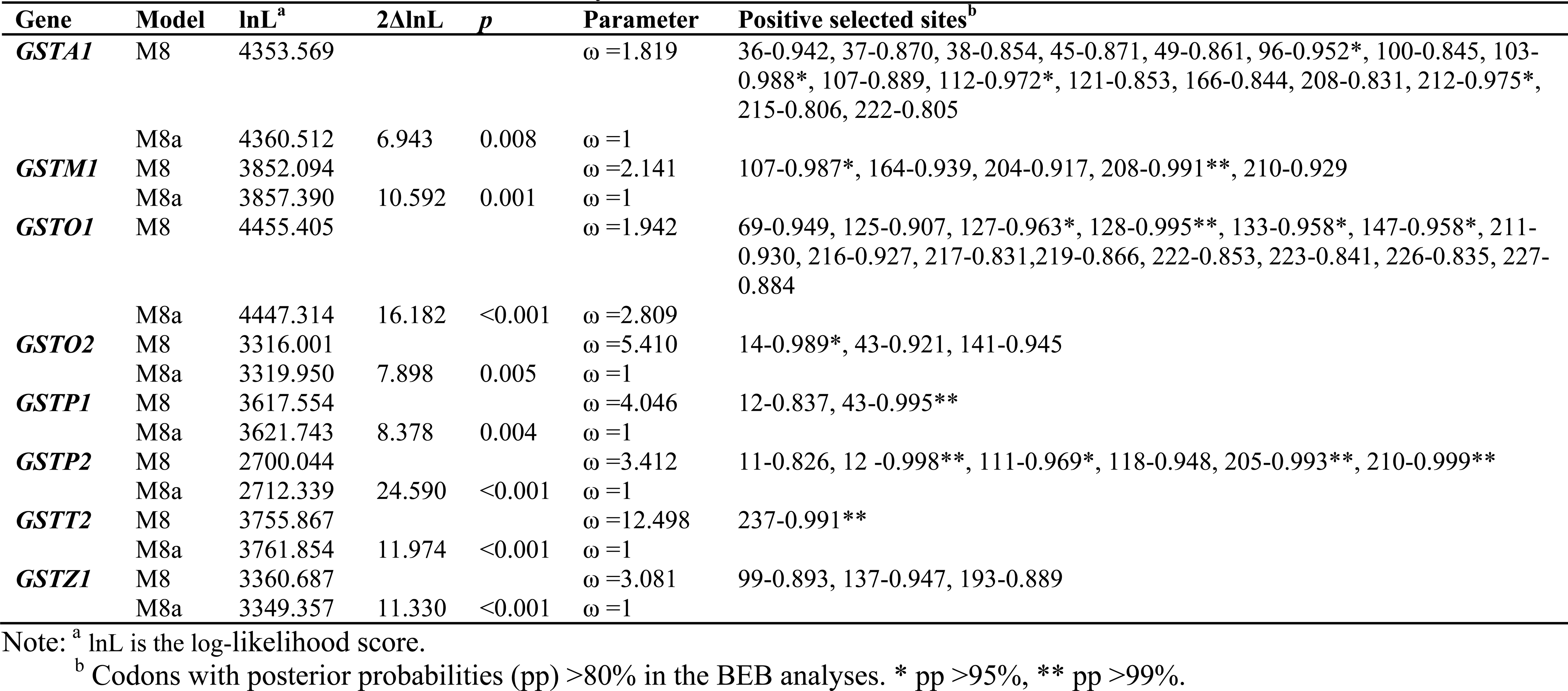
Positive selection detected in mammals by site models in PAML.

**Table S5.**
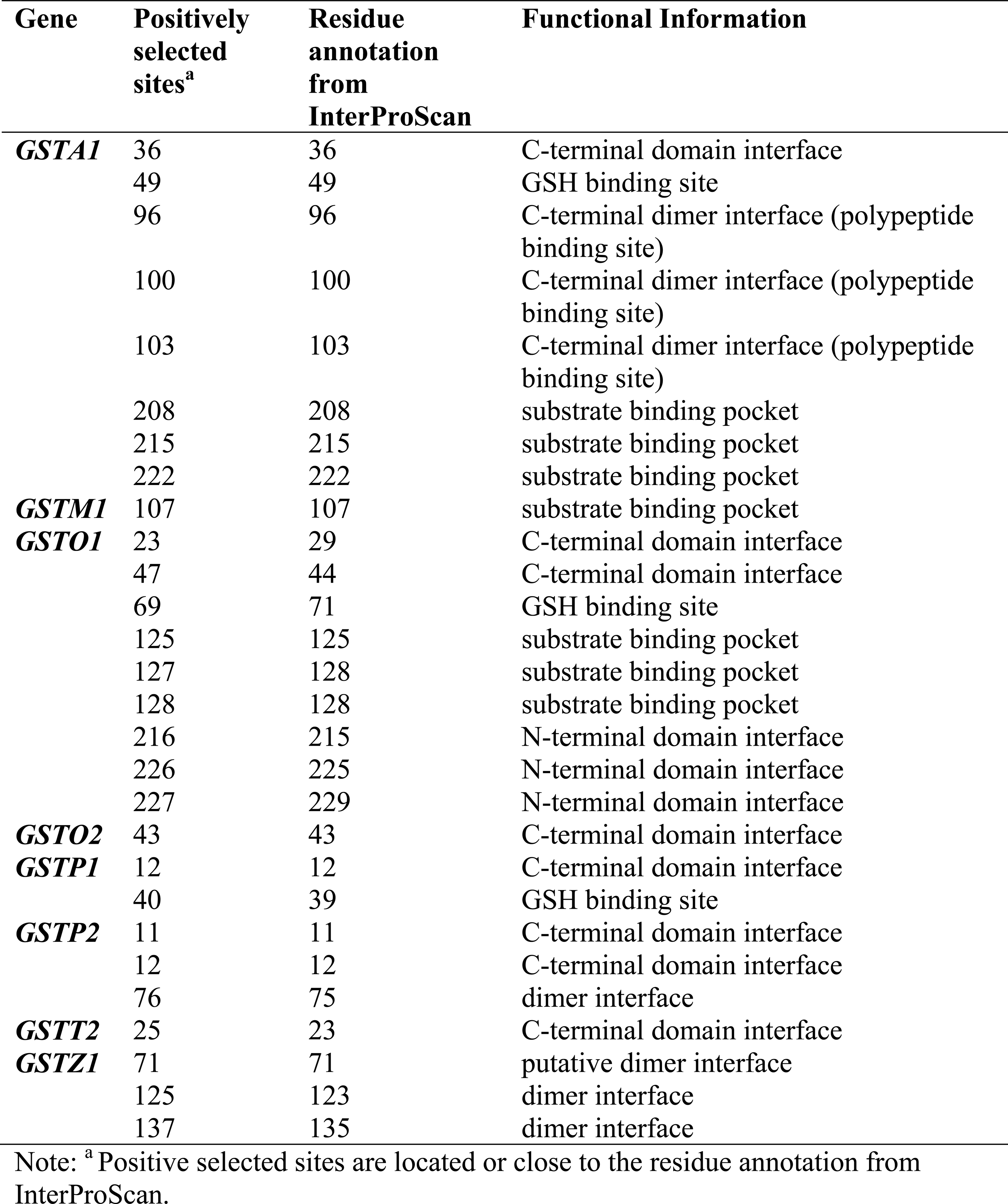
Identification of the domain location of each positively selected sites.

**Table S6.**
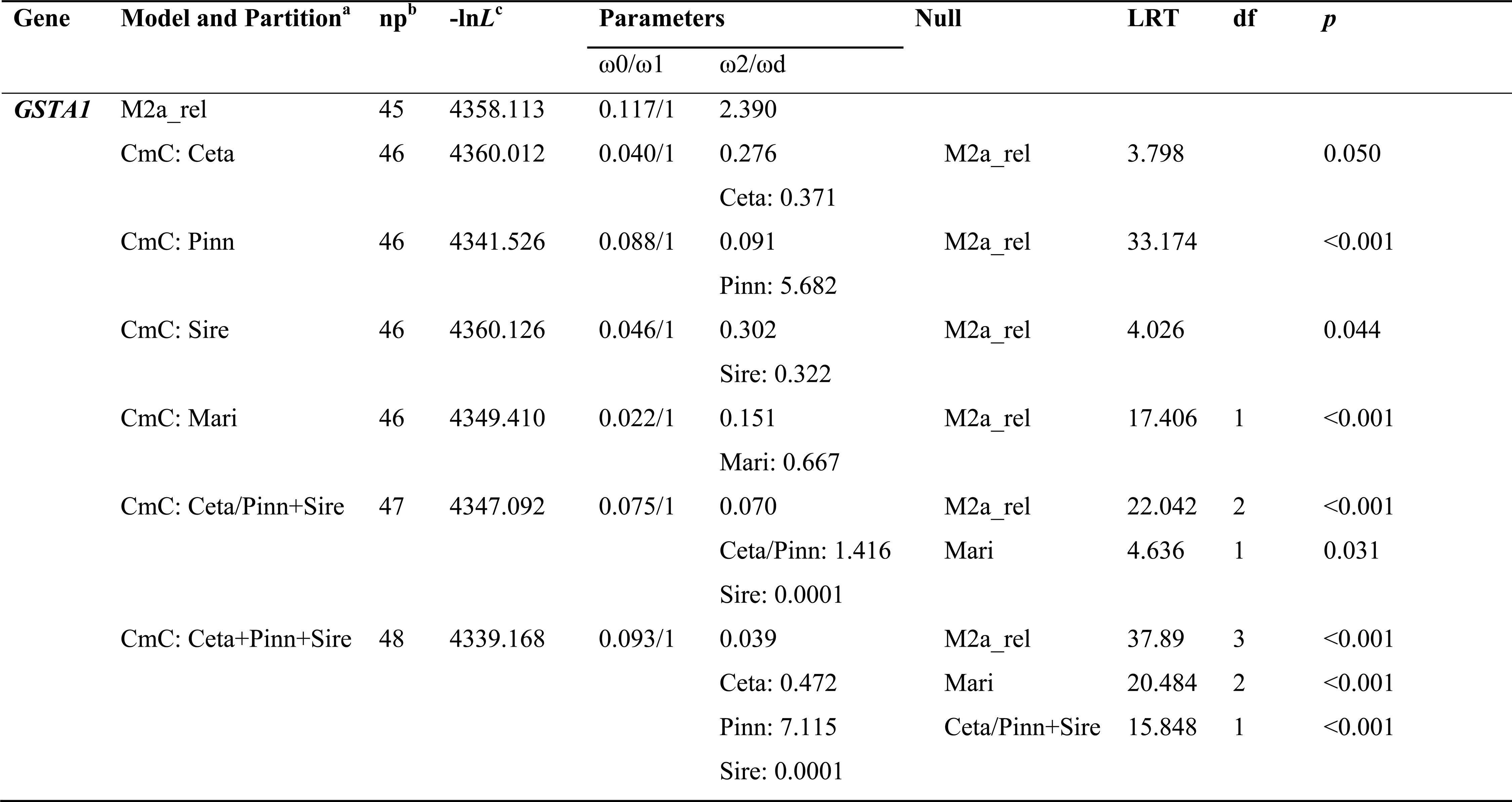
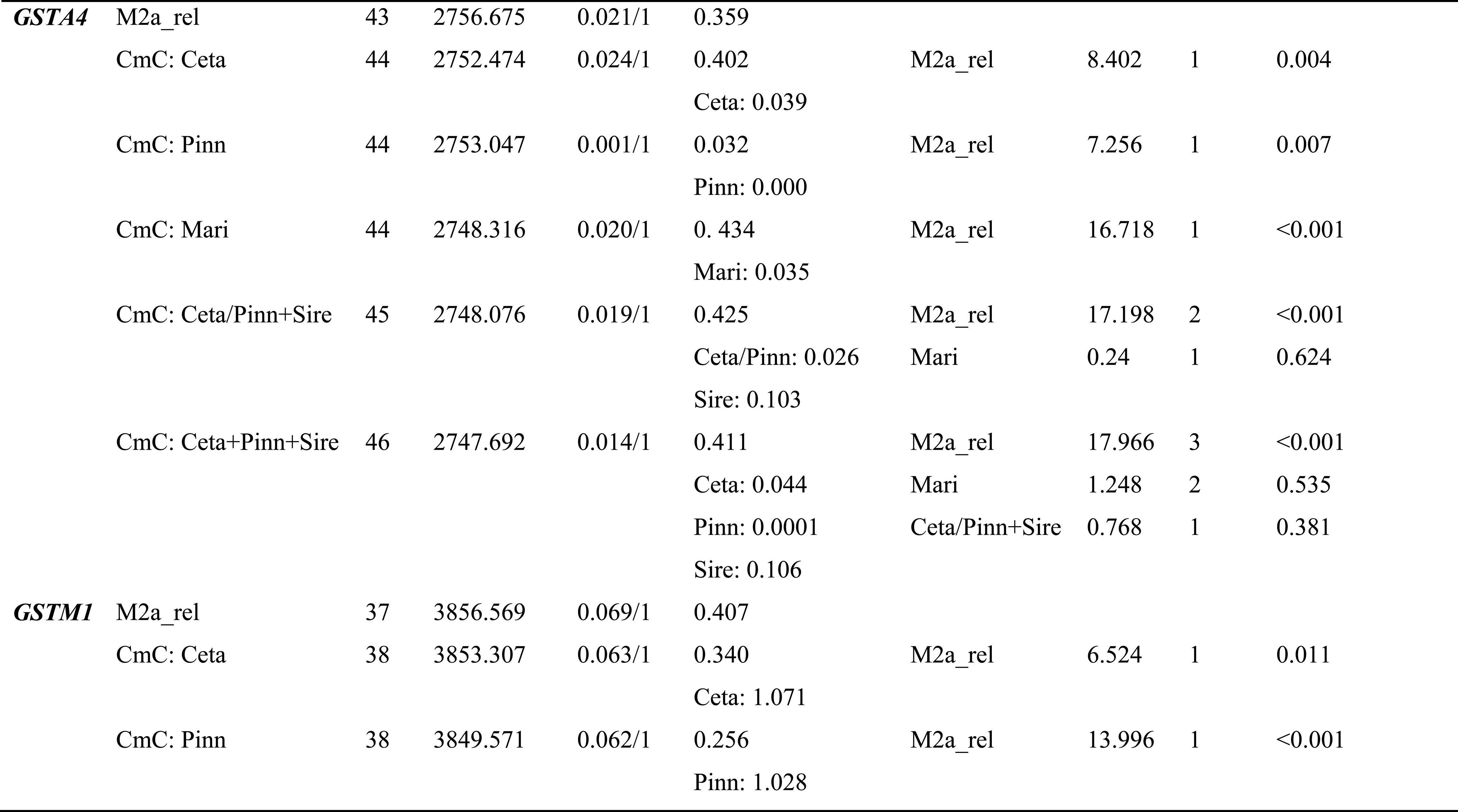
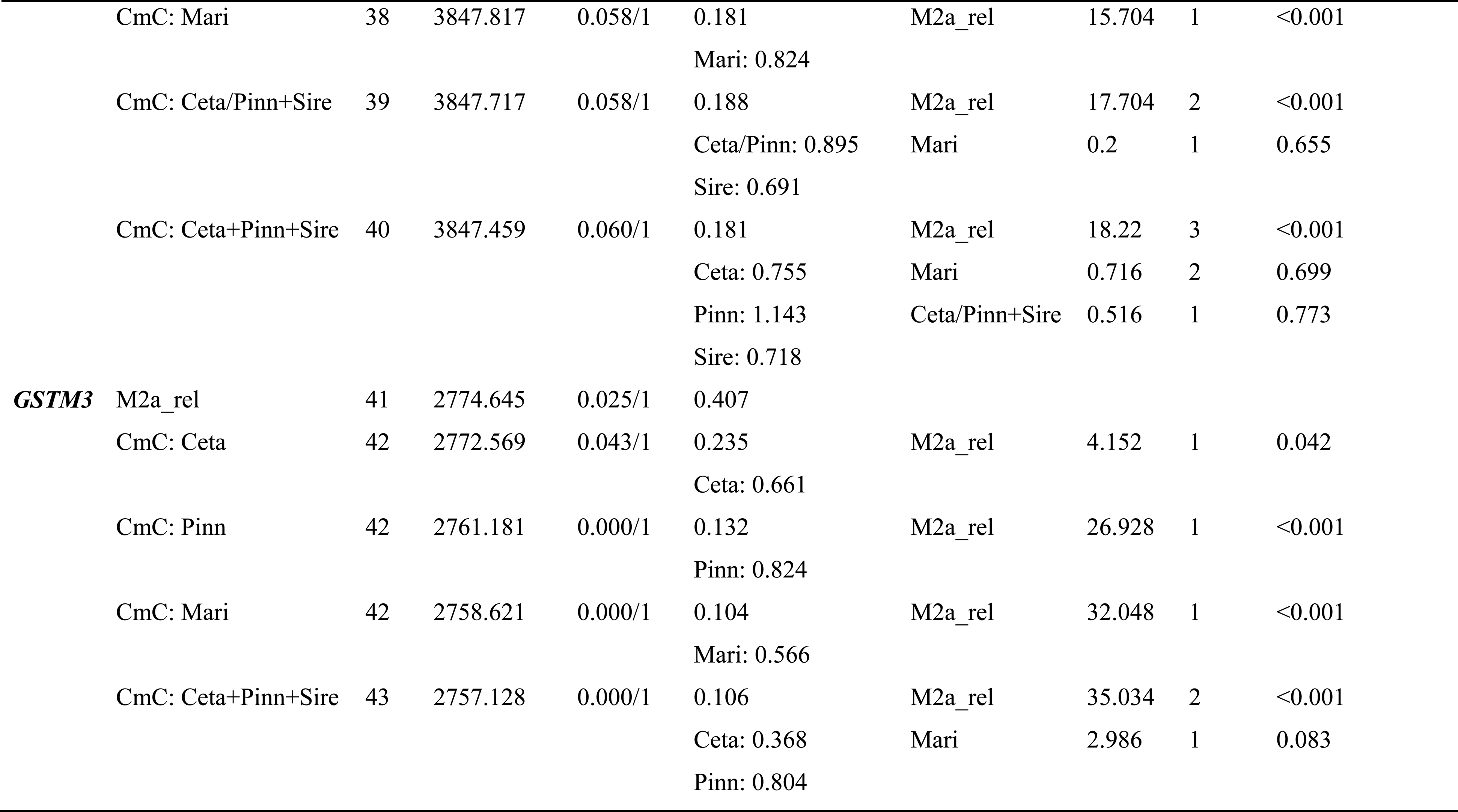
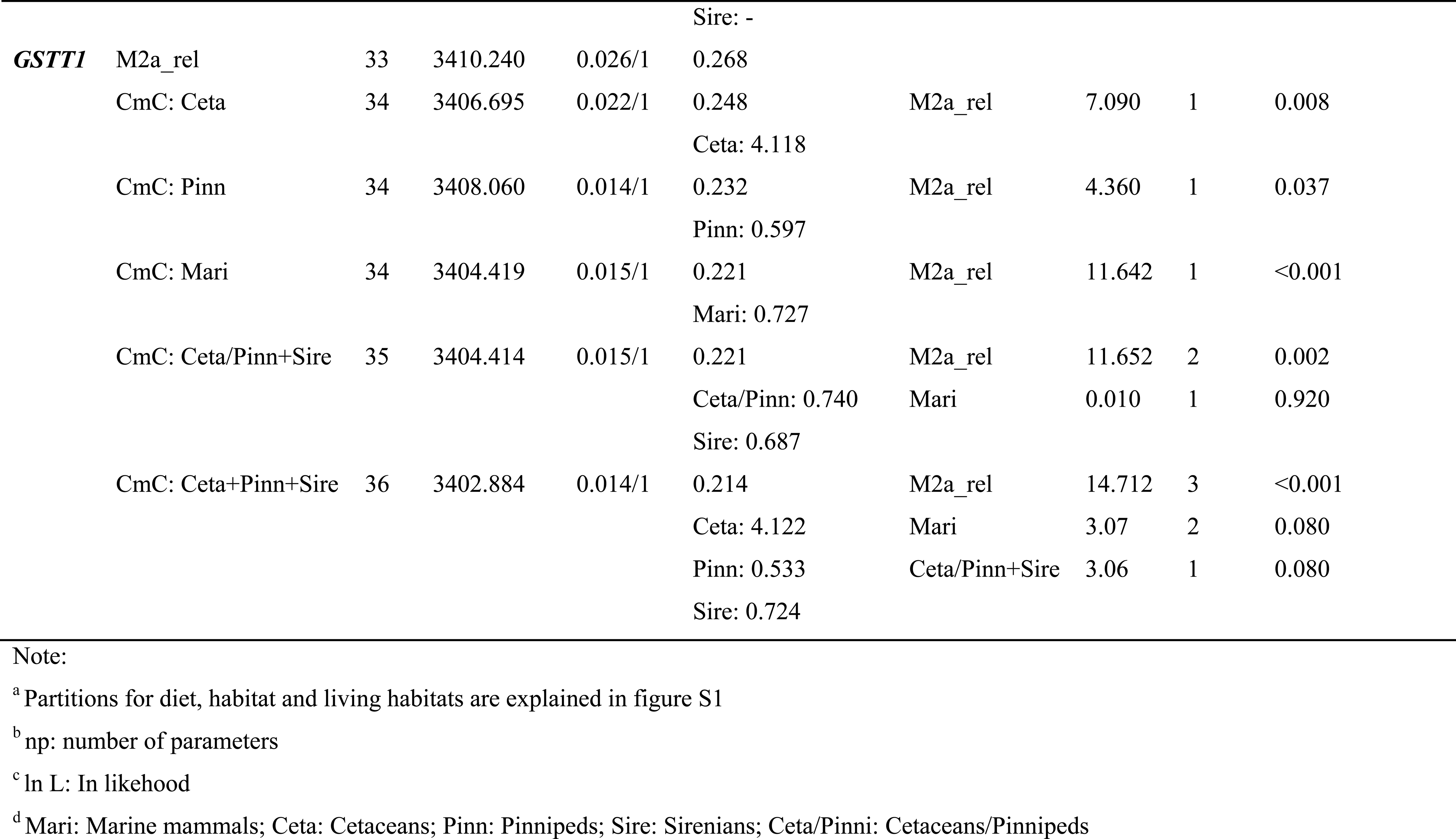
Results from Clade model C (CmC) test for divergent partitioned by habitats.

**Table S7.**
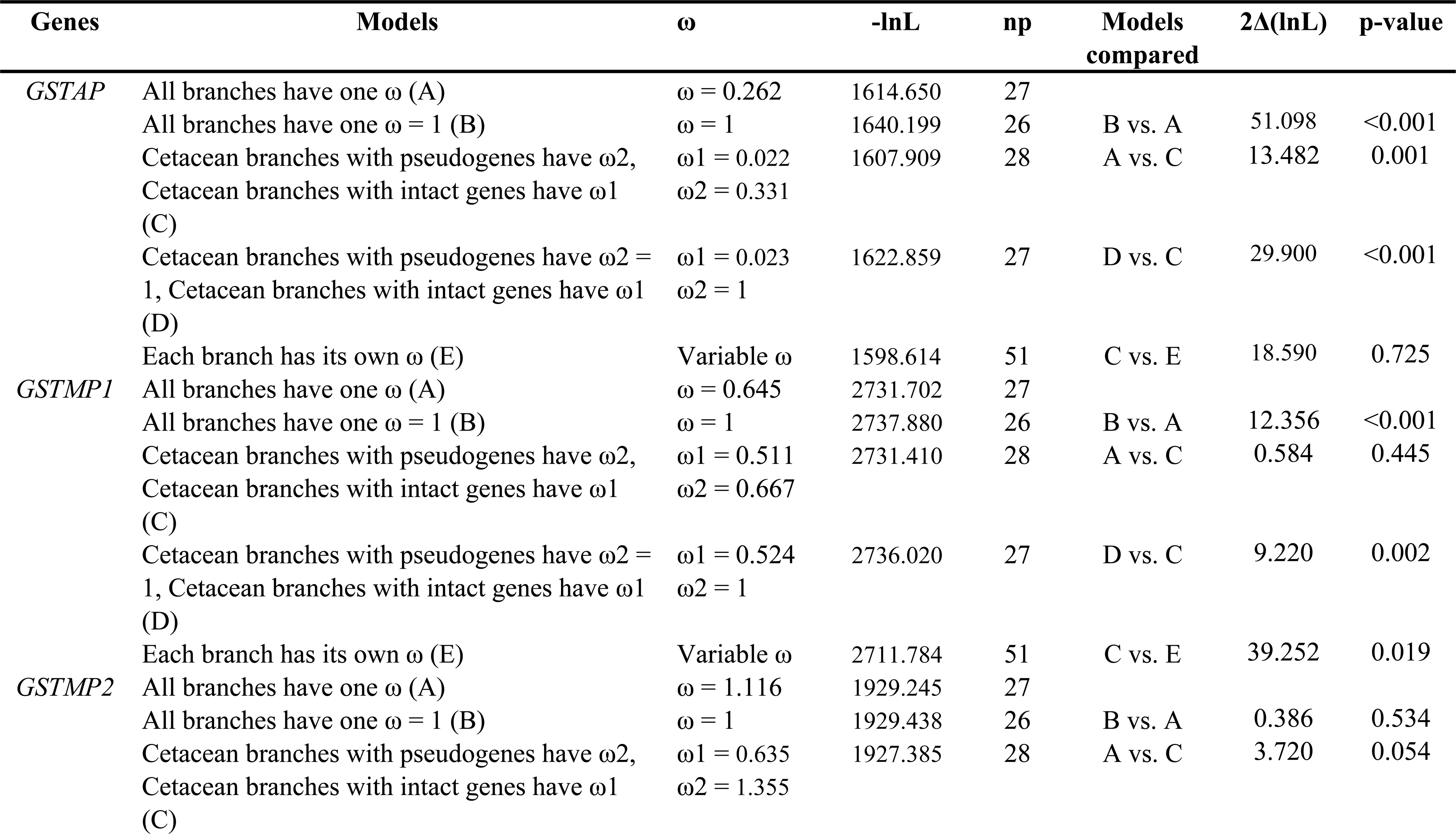
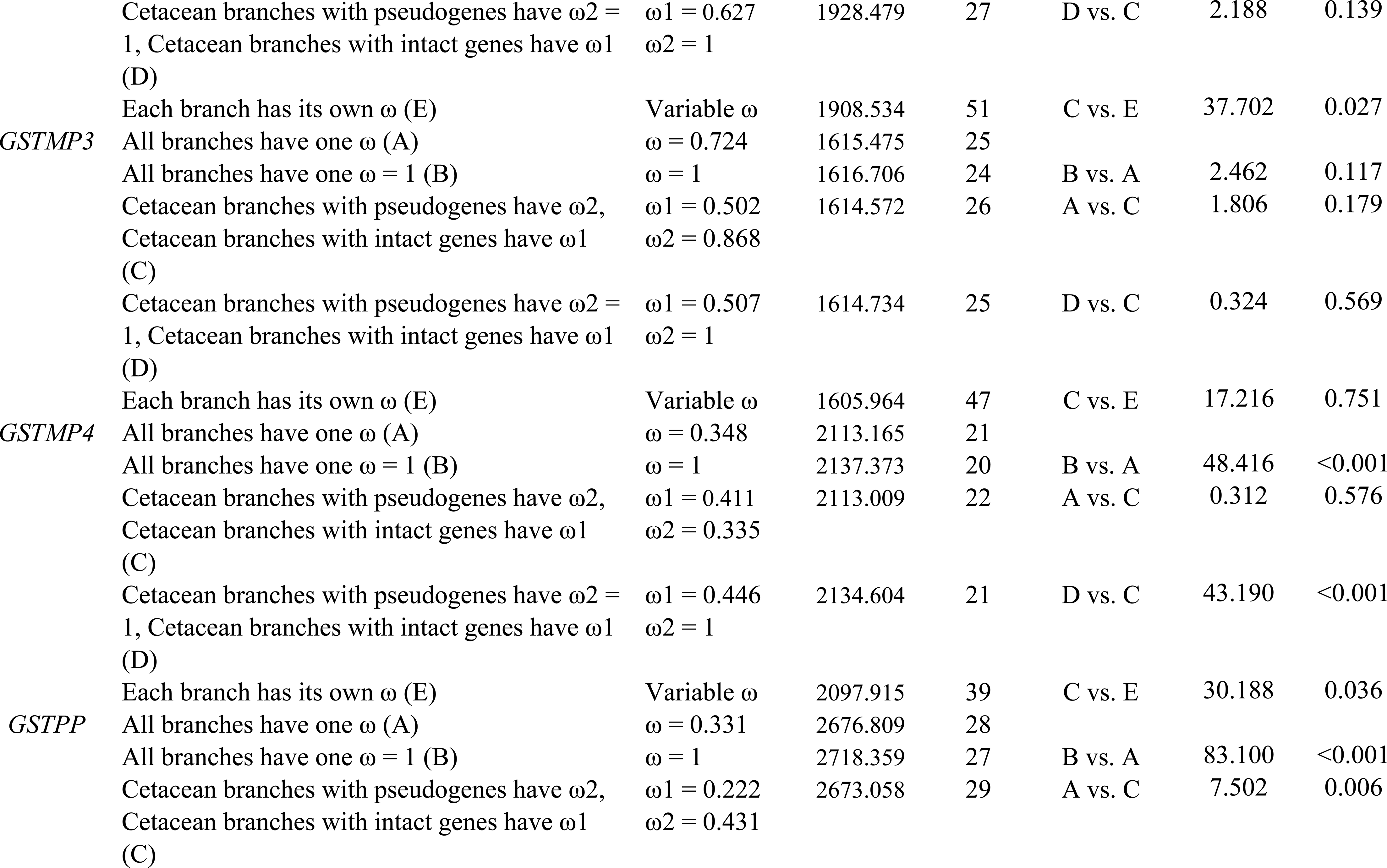
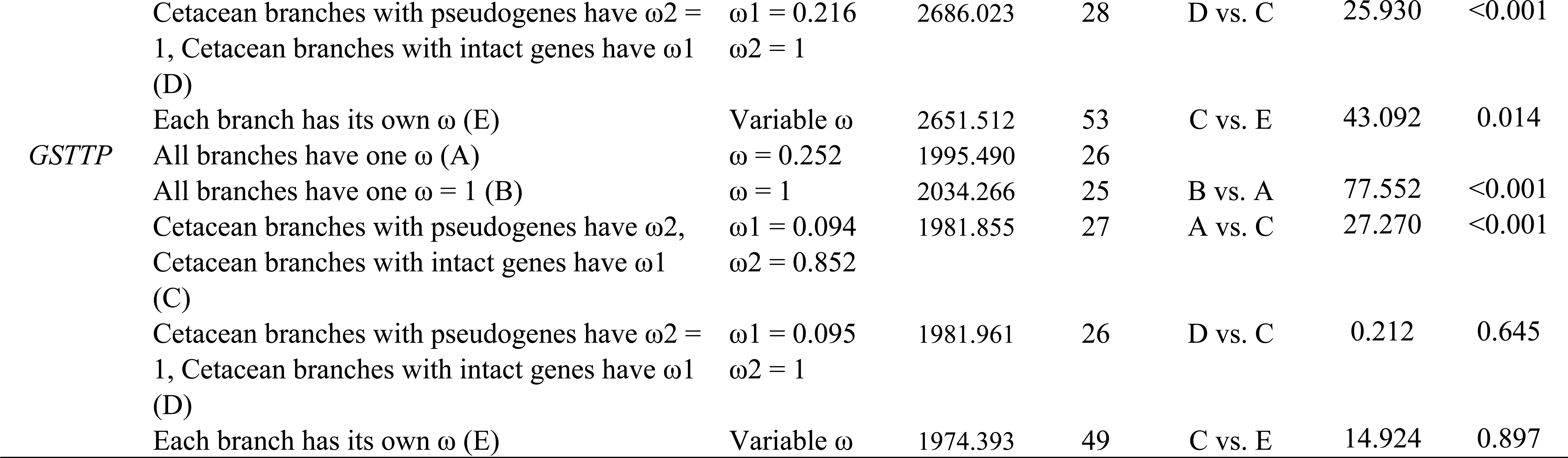
Likelihood ratio tests of various models on the selective pressures on pseudogenes in the cytosolic GST subclass in cetacean lineages.

